# Isolation, identification and selection of bacteria with the proof-of-concept for bioaugmentation of whitewater from woodfree paper mills

**DOI:** 10.1101/2020.12.14.398669

**Authors:** Nada Verdel, Tomaž Rijavec, Iaroslav Rybkin, Anja Erzin, Žiga Velišček, Albin Pintar, Aleš Lapanje

## Abstract

When whitewater circuits in the paper industry are closed, organic compounds accumulate and cause adverse production problems, such as the formation of slime and pitch. Since wood-free whitewater is usually a mixture of additives for paper production and an efficient cost-effective purification technology for their removal is lacking, the aim of our study was to find an effective bio-based strategy for whitewater treatment using a selection of indigenous bacterial isolates. We first obtained a large collection of bacterial isolates and then tested them individually for their ability to degrade the organic additives used in papermaking, i.e. carbohydrates, resin acids, alkyl ketene dimers, polyvinyl alcohol, latex, and azo and fluorescent dyes. Out of the 355 bacterial isolates, we selected a combination of four strains (*Xanthomonadales bacterium* sp. CST37-CF, *Sphingomonas sp.* BLA14-CF, *Cellulosimicrobium* sp. AKD4-BF and *Aeromonas* sp. RES19-BTP), which cover the entire spectrum of the tested organic additives. A proof-of-concept study in pilot scale was then performed by immobilizing the cells of our artificial bacterial consortium in a 33-liter tubular flow-through reactor with a retention time of <15 h. The combination of the four native strains enabled an 88% reduction in COD of whitewater even after 21 days. Additionally, we show that the bio-based whitewater treatment surpasses photolysis and photocatalysis.

**Graphical abstract:** 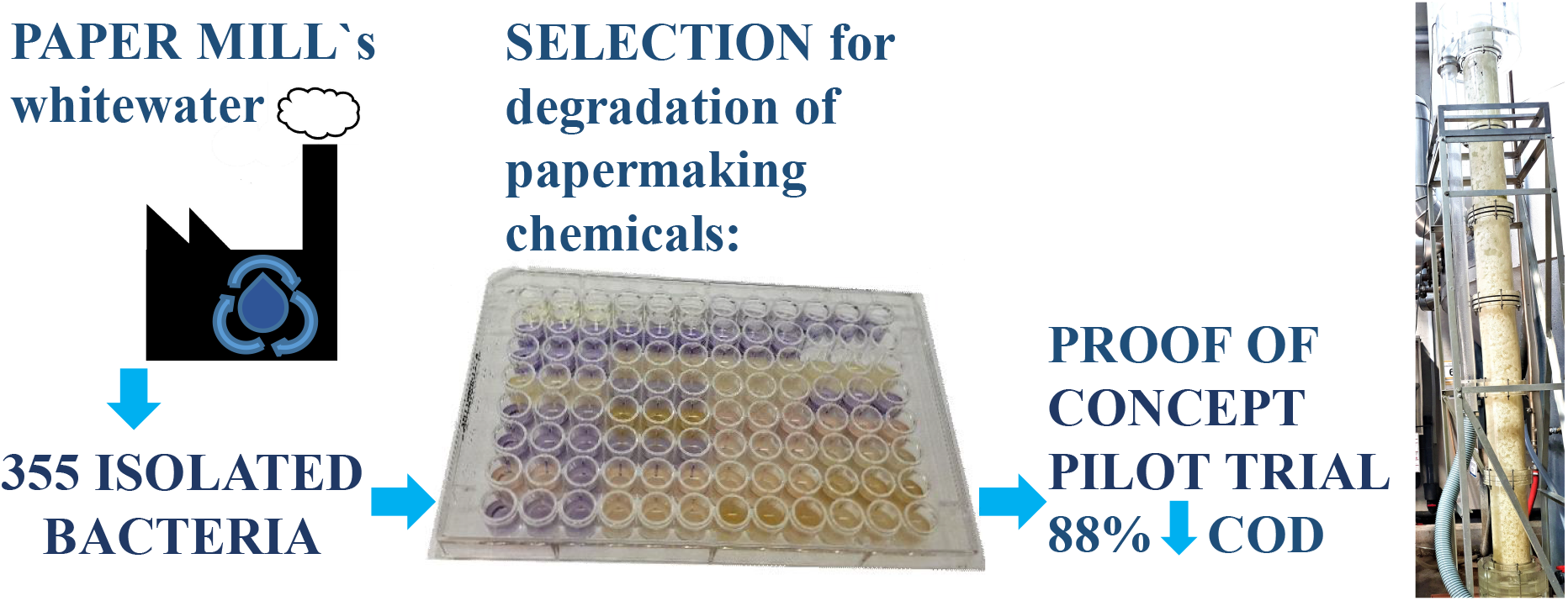

**Highlights:**

- A strategy for selecting a consortium of indigenous bacteria with a high potential for bioaugmentation of wood-free whitewater is presented.
- Study of bacterial degradation of eight chemically diverse substances used in paper production is presented. Methods for the selection of bacteria suitable for industrial application were developed.
- The constructed artificial bacterial consortium consisted of strains belonging to genera *Xanthomonadales bacterium*, *Sphingomonas*, *Aeromonas* and *Cellulosimicrobium*.
- The proof of concept of the industrial application, consisting of a 33-L-column filled with the immobilized artificial consortium, was an 88% decrease in COD of the whitewater effluent.

## 1 Introduction

The pulp and paper (P&P) industry is one of the largest consumers of water resources and causes severe water pollution. The solution currently used by the P&P industry is to reduce water consumption by applying the closed water cycle concept. However, organic compounds begin to accumulate in a closed cycle, leading to the formation of pitch and slime (Hubbe *et al.*, 2016), and new solutions to remove organic compounds from the water cycle need to be developed. In the wood-free paper industry the whitewater cycles are cleaner than in other types of paper industry because the purified imported pulp is added as a source of cellulose. Additionally, several chemicals contained in the wood-free whitewater e.g. carbohydrates, sizing agents (resin acids and alkyl ketene dimers (AKD)), binders (polyvinyl alcohol (PVA) and latex) as well as azo and fluorescent dyes, are used to ensure adequate paper quality. Despite the relatively high specific waste water volume compared to other types of paper production (m^3^ waste water per ton of paper produced) (Hamm and Schabel, 2007), to the best of our knowledge no specific treatment studies have been conducted.

The currently most common solution is based on conventional wastewater treatment with primary clarification, followed by activated sludge processes. In fact, activated sludge has proven to be effective in reducing the amount of organic compounds even at high temperatures (55°C) (Jahren *et al.*, 2002). To increase the removal of pollutants, advanced oxidation processes, anaerobic treatment and membrane filtration can be used (Hubbe *et al.*, 2016). However, they result in high sludge production, high energy consumption, low biodegradation of recalcitrant pollutants and release of secondary pollutants (Priyadarshinee *et al.*, 2016). Additionally, the choice of whitewater treatment is limited by technological parameters such as constant temperature and pH value before and after treatment (Malmqvist *et al.*, 1999). By the accumulation of organic compounds like carbohydrates, increased biological oxygen demand values lead to mucus growth in the system (Hubbe, 2007). The accumulation of sizing and binding agents causes pitch problems and impairs paper properties (Serreqi *et al.*, 2000; Miranda *et al.*, 2008). Additionally, residues of azo dyes are problematic because they are highly visible, while fluorescent stilbene dyes are recalcitrant and toxic to aquatic life (Salas *et al.*, 2019). We found that wood-free paper production systems are so specific that photolysis and photocatalysis with titanium dioxide are ineffective (Suppl. A, Section A.1). The suspended organic compounds are strongly adsorbed on the catalyst surface and block active sites on the catalyst.

The use of microorganisms can be a good alternative to other approaches because they have a low environmental impact, lower costs and the ability to degrade even the most recalcitrant organic compounds (Santisi *et al.*, 2015). In the microbial community of activated sludge, and particularly in the community established in closed whitewater circuits, it is expected that some microorganisms will be specialized in the degradation of resistant compounds and their use could be suitable for bioaugmentation. Bioaugmentation has a potential for the treatment of industrial wastewater with selected highly efficient microorganisms added to the wastewater to degrade specific pollutants or to reduce the overall chemical oxygen demand (Herrero and Stuckey, 2015). State of the art studies on the bioaugmentation of P&P industrial effluents have mainly focused on the degradation of recalcitrant lignin (Mathews *et al.*, 2014), which does not cause problems in wood-free whitewater. However, studies that are more appropriate for the wood-free industry have focused on the microbial degradation of resin acids (Yu and Mohn, 2001, 2002) and the total chemical oxygen demand of a degradation process (Li *et al.*, 2018). Ubiquitous microorganisms have been reported to hydrolyze starch (de Souza and Magalhães, 2010) and cellulose (Rabinovich *et al.*, 2002). Even the biodegradation of azo dyes has been observed on several occasions (Khan *et al*., 2013). Studies have been reported on the biodegradation of synthetic binders such as PVA (Shimao, 2001). There are still no data on good inoculums for biodegradation of AKD and fluorescent dyes.

An important problem when starting the biological treatment plants are poor start-up inoculums taken from similar environments. According to the Pareto Principle 20/80 (Dejonghe *et al.*, 2001), it is expected that only 20% of the bacteria in a microbial community perform 80% of the degradation. Moreover, degraders of recalcitrant compounds in conventional biological treatments are usually outcompeted by the faster growing degraders of readily biodegradable compounds. Here, the immobilisation of bacteria on porous carriers can protect the microbes from competitors and from fluctuating process conditions and prevent cellular washout from the bioreactors (Magrí *et al.*, 2012; Horemans *et al.*, 2017a). Therefore, it would be ideal to acquire the smallest robust consortium consisting of highly active bacteria that occupy specific ecological niches, defined by the nutrients they can degrade and utilise.

We put forward the hypothesis that bacteria capable of decomposing chemicals for paper production are found in the whitewater itself. Probably most of them are adapted to decompose the most readily degradable carbon sources, such as starch, while others are able to decompose the less biodegradable compounds, even fluorescent dyes. Complex chemical mixtures require either the cooperation of several specialists that complement each other in different niches or a single generalist with versatile metabolic processes. However, consortia of bacteria are more advantageous for handling different nutrient concentrations than a single strain (Santisi *et al.*, 2015).

Accordingly, the main aim of this study was to simulate the real system and artificially construct bacterial consortia capable of degrading all the chemicals introduced in the wood-free paper manufacturing process (starch, cellulose, resin acids, AKD, polyvinyl alcohol, latex and azo and fluorescent dyes). In preparing the consortium, we pursued specific aims (Fig. 1): (i) Isolation of individual strains capable of degrading a range of papermaking chemicals; to isolate bacteria with stable catabolism at low nutrient concentrations, we performed bacterial isolation under nutrient-poor conditions (Horemans *et al.*, 2017b), (ii) construction of the most suitable inoculum; we prepared different combinations of active bacteria in synthetic and real whitewater to select the combination with the broadest repertoire of carbon source use, and (iii) upscaling the use of the prepared inoculum by applying an immobilisation strategy.

**Fig. 1.**
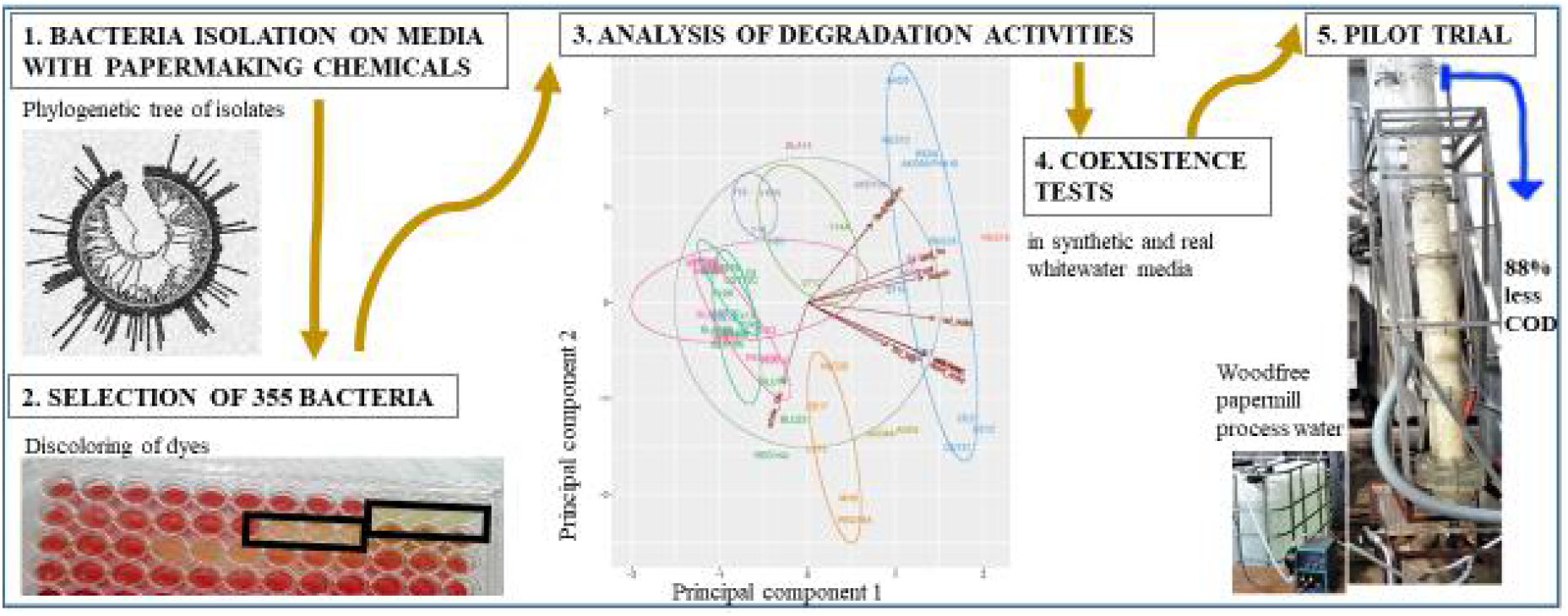
Scheme of the strategy for construction of a native bacterial consortium for bioaugmentation of wood-free whitewater.

## 2 Methods

### 2.1 Whitewater analysis and initial treatment

The whitewater from a wood-free paper mill was systematically analyzed. Over a period of two months the chemical oxygen demand, COD (ISO 6060), biological oxygen demand, SBOD (ISO 5815-1), dissolved organic carbon content (DOC) (Teledyne Tekmar, model Torch analyzer) and pH value (Metrohm, model 827) were measured. The cation content (Ca^2+^, Mg^2+^, Na^+^, K^+^, NH_4_^+^) was determined with an ion chromatograph (Dionex Thermo Fisher Scientific) and the concentrations of anions (Cl^−^, SO_4_^2−^) were measured with a Dionex DX-120 ion chromatograph. The content of HCO_3_^−^ ions in the investigated samples was titrated with HCl (see results in Supplementary Materials A, Table A.1). Initial unsuccessful treatment results with photocatalysis and photolysis are described in Suppl. A (Section A.1).

### 2.2 Sample collection

The wood-free paper mill produces 87,000 tons of wood-free specialty, coated and uncoated paper per year. The whitewater circuits consist of a closed short and a long circuit. In the long circulation, the whitewater flows through the disc filter, where it is clarified in countercurrent through a pulp fiber cake. It is discharged as clear filtrate (CF), which is reused for dilution in stock preparation. At regular intervals CF is discharged into a biological treatment plant from where it is discharged into surface waters. CF, biofilm from the wet-end (BF) and the effluent from the biological treatment plant (BTP) were individually microbiologically enriched in M9 media (VWR, USA) (Miller, 1972) and used as a source of the collected bacterial strains. After 7 days at 25°C 100 μL of the 100-fold diluted grown cultures were plated in triplicates onto corresponding solid M9 media. The M9 media was supplemented with 0.3 mM papermaking additives as the only carbon source. The following carbon sources were used: (i) readily biodegradable compounds (AKD, resin acids, cationic, native and soluble starch, cellulose, PVA and latex), (ii) none-readily biodegradable azo dyes (red, blue, black and yellow), and (iii) fluorescent whitener. In this way, we isolated bacteria that are capable of separately using 13 different sources of carbon. The isolates were marked according to the carbon source, as in Suppl. A, Table A.2. For example, isolates from media with AKD as the only carbon source (M9 AKD) were marked sequentially as AKD1, AKD2, etc. The list of bacterial isolates stored in 25-30% glycerol solutions is given in Supplementary Material B, Table B.1.

### 2.3 Biodegradation assays

Cycloheximide in a concentration of 10 mg/L (Sigma Aldrich, USA) was present in all biodegradability tests. Using microplate assays, we determined growth, absorption or fluorescence of carbon sources with a microplate reader (Synergy H4 (Biotek, USA)). We used untreated, sterile 96-well flat-bottom microplates (VWR, USA) to incubate the bacteria. We incubated 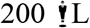 media with and without bacteria at 25°C and 50 rpm. Using calibration curves, we calculated the coefficient between the unused and the initial concentration of the carbon source, C/C_0_.

#### 2.3.1 Readily biodegradable compounds

For the selection of the degraders of readily biodegradable compounds, similar methods were applied as described previously for starch by Lal and Cheeptham (2012), with modifications. In short, 0.25-10 g/L starch, cellulose, AKD, resin acids, PVA or latex were added to the M9 minimal agar media as sole carbon sources. We inoculated four evenly spaced 0.8 cm^2^ circular areas per 41 cm^2^ surface area of the Petri dish, as in Suppl. A, Fig. A.1e, and incubated for up to 8 days at 25°C. After incubation, cell viability was checked on solid NB medium. Iodine solution (Lugol, I_2_+ KI; Merck, Sigma Aldrich) was then poured onto the plates. Where a discoloration ring around active colonies was detected, the ring radius was measured. The percentage of unused carbon source was calculated using Equation A.1 (Suppl. A, Section A.2). Degraders of cationic and soluble starch and AKD were additionally screened in liquid media. Absorbance measurements at 584 nm, before (C_CELLS_) and after (C) addition of Lugol were recorded. Since cell removal was not possible due to the non-Newtonian properties of the media, the (C-C_CELLS_)/C_0_ values were compared; C_0_ marked the initial concentration of the carbon source.

#### 2.3.2 Azo dyes

The decoloration experiments with the red, blue and black azo dyes were similar to those described previously (Parmar and Shukla, 2018), with modifications. Bacterial activities were determined in M9 media containing 47-95 mg/L dye as sole carbon source. Cometabolism, which we refer to simultaneous biodegradation of two substances, of azo dyes was tested in NB or M9 glucose containing the dye (5-10 mg/L). For the cometabolism tests, bacteria were incubated in media until the first discoloration, which occurred after 18-72 h. Before determination of the discoloration, the samples were centrifuged at 12,000g for 5 minutes. The supernatant was used to measure the discoloration at the specific Abs maxima of the dye (Suppl. A, Table A.2). Due to sedimentation of the yellow dye particles, which prevented removal of the cells by centrifugation, we used two protocols for yellow dye degradation (Suppl. A, Section A.2.2). The first protocol was similar to that described previously (Parmar and Shukla, 2018), with modifications. Bacteria grew in NB media. The four fastest growing isolates were cleared in 9 g/L NaCl and incubated in M9 with 37 mg/L yellow and after 48 h Abs-maximas were recorded (400 nm) and cfu counted. For the second protocol, a bacterial colony was picked and transferred to mineral media supplemented with 82 mg/L yellow (C) or 47 mg/L glucose. Incubation in mineral media with 47 mg/L glucose was used to reflect the absorption of the cells (C_CELLS_). After 168 hours of gentle shaking at less than 50 rpm and room temperature, the absorbance was recorded at 400 and 600 nm.

#### 2.3.3 Fluorescent whitener

The biodegradation tests were performed in M9 glucose containing whitener (5-900 mg/L). Precise concentration values for each test separately are given in Suppl. A, Table A.3. To determine the fluorescence intensity of whitener after excitation at 315 nm, the emission maximum was measured at 430 nm. The bacteria were incubated until OD_600_ increased significantly, i.e. up to four times. Before determining the degradation of whitener, bacteria were removed from media by centrifugation at 12,000g for 5 min and the supernatants were transferred to black flat-bottomed microtiter plates (VWR).

### 2.4 Selection of bacteria

To find bacterial isolates with the highest bioaugmentation potential, we selected them according to the following protocols (Suppl. A, Table A.3):

(i) Out of 355 bacteria, we selected 2-5 of the most active bacteria per carbon source used to isolate the respective bacteria, i.e. 47 isolates (Suppl. A, Fig. A.1). For the inoculum, a colony of the isolate was transferred or smeared by a sterile loop. The most active bacteria were those that degraded (or used) the most of carbon source for a given test period. Additionally, the time-dependent activity of the most active degraders of blue, red and whitener was determined. For the inoculum, the isolates were incubated for 3 days at 25°C in a shaking incubator at 50 rpm in a suitable M9 medium. 10-20 μL of the culture was transferred to NB medium containing a suitable dye or a glucose-enriched M9 whitener. The use of the carbon source was determined at certain intervals.
(ii) The selected 47 bacterial isolates plus 15 primary and secondary contaminants were further tested for their repertoire of carbon sources (specialists vs. generalists) (Suppl. B, Table B.2). For inoculum, the isolates were cultured in glucose M9 medium for 3 days at 25°C. We tested bacteria for the use of the following carbon sources: cationic and soluble starch, cellulose, resin acids and AKD, azo dyes (red and blue) and whitener (Suppl. A, Fig. A.2 and Suppl. B, Table B.3). Furthermore, the growth of the 6 selected isolates was calibrated by determining OD_600_ vs. cfu (Fig. A.3).
(iii) The coexistence of the 6 selected bacteria was tested for all 63 combinations. We selected a combination of bacteria that utilizes the most organic carbon content in sterile M9 media with real and synthetic whitewater after a certain time. To create similar conditions in both media, synthetic media was composed of CST, PVA, resin and direct blue 15 dye in the respective weight ratios 1:1:1:0.5. Real whitewater was sterilely filtered through 0.2 mm membrane PP filter (Macherey-Nagel) before the experiments. For inoculum, the individual overnight cultures of the six bacteria in NB were harvested and washed three times in 9 g/L NaCl. OD_600_ was recorded and cfu calculated. The combinations of bacteria had the same initial sum of evenly distributed bacteria with cfu 10^7^/mL. The experiment was performed in six replicates in 96-well microtiter plates. After 3 days at 25°C the media were sterilely filtered through 0.2 mm PP filters. The complete mineralization of the organic content was analyzed by DOC measurements with a TOC analyzer (Shimadzu).

### 2.5 Pilot-scale experiment (the proof of concept)

For proof of concept, the consortium of four isolates was immobilized on carriers and placed in a 33-liter column with unsterile whitewater. A continuous parallel flow of 2 L/h whitewater and air through the column (r=0.06 m, h=2 m) was introduced at the bottom by a Watson Marlow pump. The hydraulic retention time in the column was <15 h. There was almost no mixing of the carriers and the whitewater was saturated with oxygen. To achieve the ratio of COD:N:P = 100:5:1, which is necessary for the growth of bacteria, urea and H_3_PO_4_ were added to the original whitewater. Immobilization of the isolates on carriers was performed as reported (Horemans *et al.*, 2017a). Inoculum was prepared as follows: The 4 isolates were separately grown 48 h at 25°C NB media. To the harvested bacteria (15 minutes at 12000g) 0.5% sodium alginate and sterile carriers Kaldnes K3 (Tongxiang Small Boss Ltd., China) were added. Initially, cfu/mL of RES19, CST37 and BLA14 on immobilized supports was 7×10^5^±16% and of AKD4 4×10^6^ cfu/mL. The mixture was vacuumed three times with a vacuum pump ILMVAC. 1.6 L of carriers loaded with bacteria, 0.4 L for each isolate, were placed in a container containing approximately 25 L of carriers. The next day they were placed in the column. An oxygen electrode (Hach) was positioned at the top of the column. The following process parameters were measured with rapid tests (Nanocolor): total N, total P, PO_4_^3−^, NO_3_^−^, NH_4_^+^ and COD. The pH value was measured with a portable pH meter (Hach). The seventh day, cfu/mL on carriers, taken at 20-30 cm below the top of the column, was counted as follows: the carriers were placed in 50 mL sterile centrifuge tubes containing 20 mL sterile 9 g/L NaCl (4 carriers per tube). The contents was homogenized 5 minutes at maximum vortex speed (IKA genius 3), serially diluted to 10^−10^ in 9 g/L NaCl. 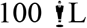 of the dilutions were spread on NB agar media and incubated at 37°C for 48 hours. Due to differences in the morphology of the bacterial colonies, we counted cfu separately for each isolate by Motic SMZ-168 microscope. After eight days an additional Watson Marlow qdos 30d pump was added to establish a recycle with 2 L/h. The experiment was followed until the 21st day.

### 2.6 Identification

DNA of 123 bacterial isolates was obtained using Chelex®100 Resin (Walsh *et al.*, 1991). One colony of defrozen isolated bacteria on NB was transferred into the chelex solution. After 15 minutes at 95°C and 15 minutes 5000 rpm centrifugation at 4°C the supernatant was transferred into RNA-free tubes and stored at −20°C. 16S rRNA gene was PCR amplified using standard primers according to (Hubad and Lapanje, 2013), with modification. 27fwd (AGA GTT TGA TCM TGG CTC AG) and 1492rev (CGG TTA CCT TGT TAC GAC TT) (Integrated DNA Technologies, USA) primers were used.

In order to check the PCR reaction, PCR samples with a 1 kb DNA ladder, DNA gel loading dye and SybrGreen I, a fluorescent nucleic acid stain, were loaded for electrophoresis. Products of the amplification were inspected on the 1% agarose gel. All PCR products were Sanger sequenced with forward primer at Macrogen, NL.

### 2.7 Phylogenetic trees

Before any further bioinformatic procedures, the chromatograms of 16S rRNA sequences were visually inspected in Chromas 2.6.6 software developed by Technelysium Pty. Ltd. Incorrect parts and ambiguous bases to the degenerate code according to the IUPAC nomenclature were deleted or modified. Finally, 1090-1340 bp large sequences were obtained. The ten most similar sequences to the sequences of our isolates were obtained by the BLASTN 2.9.0+ software. Data was retrieved on the RDP (Ribosomal Database Project) and NCBI (National Center for Biotechnology Information). The modified sequences were deposited in the NCBI GenBank with accession numbers MW144826 - MW144947 and MW131116.

Two types of phylogenetic trees were created on the basis of the Kimura 2-parametric model in MEGA X software (Kumar *et al.*, 2018). The clustering of 122 isolates together with the most similar type strains was inferred using the unweighted pair group method with arithmetic mean (UPGMA). The evolutionary distances were computed using the p-distance method, whereas the clustering of the four isolates of consortium together with the most similar type strains was conducted by using the NJ algorithm on the distance matrix.

### 2.8 Principal component analysis

Statistical correlation between carbon source utilization, genera and inoculum site of the isolates was studied by principal component analysis (PCA). The degradation was expressed in relative units on a scale from 0 to 1. The relative utilised and initial concentration of carbon source at the end of the experiment, C_U_/C_0_, was compared. PCA was performed using R software version 3.3.3 (R Core Team, 2017). Incomplete data was imputed by the missMDA of the factominer package, PCA was calculated by the native prcomp package and the data was visualized by the ggbiplot package based on the ggplot2 package (Wickham, 2016).

## 3 Results

### 3.1 Whitewater analysis

A systematic analysis of whitewater from the wood-free paper mill over a period of two months revealed relatively constant (±10%) values of 300, 165 and 95 mg/L for COD, BOD_5_ and DOC, respectively (Suppl. A, Table A.1). Our preliminary approach to degrade these organic compounds in whitewater by photocatalysis and photolysis was not successful (Supplementary Material A, Section A.1). We were only able to degrade 1% of the organic carbon content. Consequently, we prepared an alternative solution using indigenous bacterial cells that are efficient degraders of organic additives used in the paper production process.

### 3.2 Isolation and phylogeny of native bacteria

We isolated 355 bacteria from three sources. We then identified 122 isolates, including 58 strains that were studied for the repertoire of carbon source use with PCA and most isolates from recalcitrant media, like red, blue and whitener. The ten closest relatives of the identified strains belonged to four phyla, namely, Proteobacteria, Firmicutes, Actinobacteria and Bacteroidetes (Fig. 2 and Suppl. B, Table B.4). 16 strains were identified with the closest relative [*Pseudomonas*] *boreopolis* (T), with the brackets indicating their provisional phylogenetic classification.

**Fig. 2.**
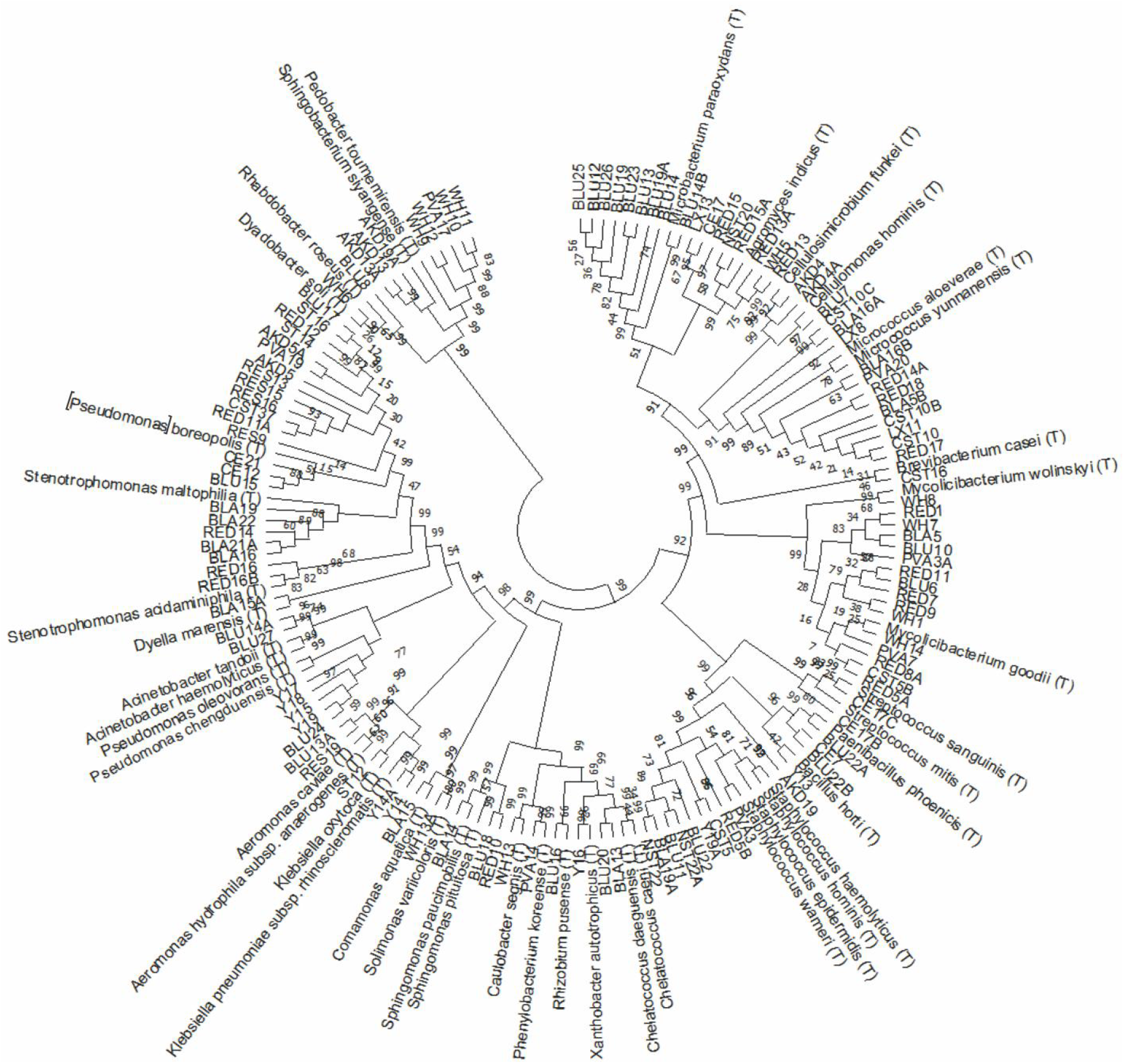
Phylogenetic tree UPGMA based on 16S rRNA sequences of 122 isolates with their closest type strains. Values at the bifurcations are showing bootstrap values of 500 clustering replications.

The highest proportion of identified isolates was obtained from CF (48%), while 26%±1% came from BF and BTP. Only ten Staphylococcus strains were isolated in all three inoculum sites, while the remaining strains belonging to the same genus were mostly derived from one inoculum site (Table 1).

**Table 1.** Distribution of genera of sequenced isolates among the inoculation sites (clear filtrate (CF), biofilm (BF) and wastewater from the biological treatment plant (BTP)); only genera with more than one strain are listed.

| Genus | Number of sequenced*<br>isolates (/) | Distribution of isolates (%) |  |  |
| --- | --- | --- | --- | --- |
|  |  | CF | BF | BTP |
| *Sphingomonas* | 3 | 100 | 0 | 0 |
| *Pedobacter* | 5 | 100 | 0 | 0 |
| *Chelatococcus* | 2 | 100 | 0 | 0 |
| *Xanthomonadales bacterium* | 16 | 90 | 10 | 0 |
| *Mycobacterium* | 16 | 90 | 0 | 10 |
| *Micrococcus* | 13 | 62 | 23 | 15 |
| *Cellulosimicrobium* | 2 | 0 | 100 | 0 |
| *Pseudomonas* | 5 | 0 | 100 | 0 |
| *Streptococcus* | 2 | 0 | 100 | 0 |
| *Stenotrophomonas* | 7 | 0 | 90 | 10 |
| *Agromyces* | 8 | 0 | 90 | 10 |
| *Microbacterium* | 9 | 0 | 0 | 100 |
| *Aeromonas* | 2 | 0 | 0 | 100 |
| *Acinetobacter* | 2 | 0 | 0 | 100 |
| *Sphingobacterium* | 3 | 0 | 0 | 100 |
| *Klebsiella* | 3 | 30 | 0 | 70 |
| *Paenibacillus* | 3 | 30 | 0 | 70 |
| *Staphylococcus* | 10 | 30 | 30 | 40 |

\*Isolates were identified by 16S rRNA sequencing.

### 3.3 Selection of bacteria

We tested bacteria using methods based on clear and precise results of visual discoloring, UV-Vis or fluorescence spectrophotometry. Of the 355 isolates, we selected 47 for their ability to degrade a particular nutrient source (Suppl. A, Table A.4, Fig. A.1). For the selection of AKD degraders, AKD-iodine complex determination was confirmed by DOC measurements (Suppl. A, Table A.5). According to our results, dye degradation could not be followed in any of the strains when dye was the only carbon source (Suppl. A, Section A.3). However, the addition of additional carbon sources to the media made it possible to test the degradation of the isolated strains for red, blue, black and whitener dye (RED, BLU, BLA and WH), Suppl. A, Fig. A.1. For the best degrading strains for RED, BLU and WH, with the highest methodological accuracy, we determined the time-dependent activity for the respective dyes for up to 168 hours (Fig. 3 and Suppl. A, Table A.6).

**Fig. 3.**
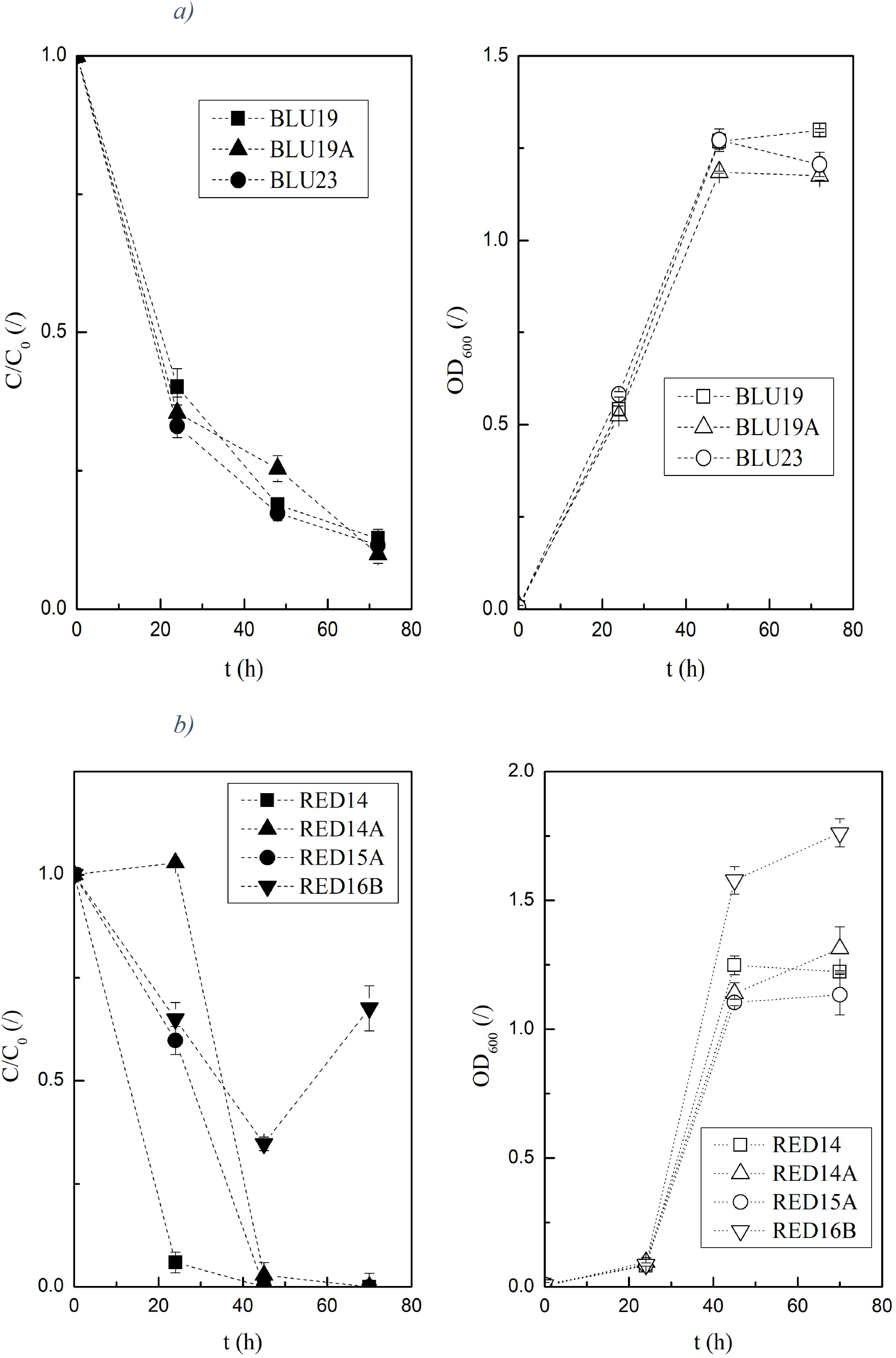
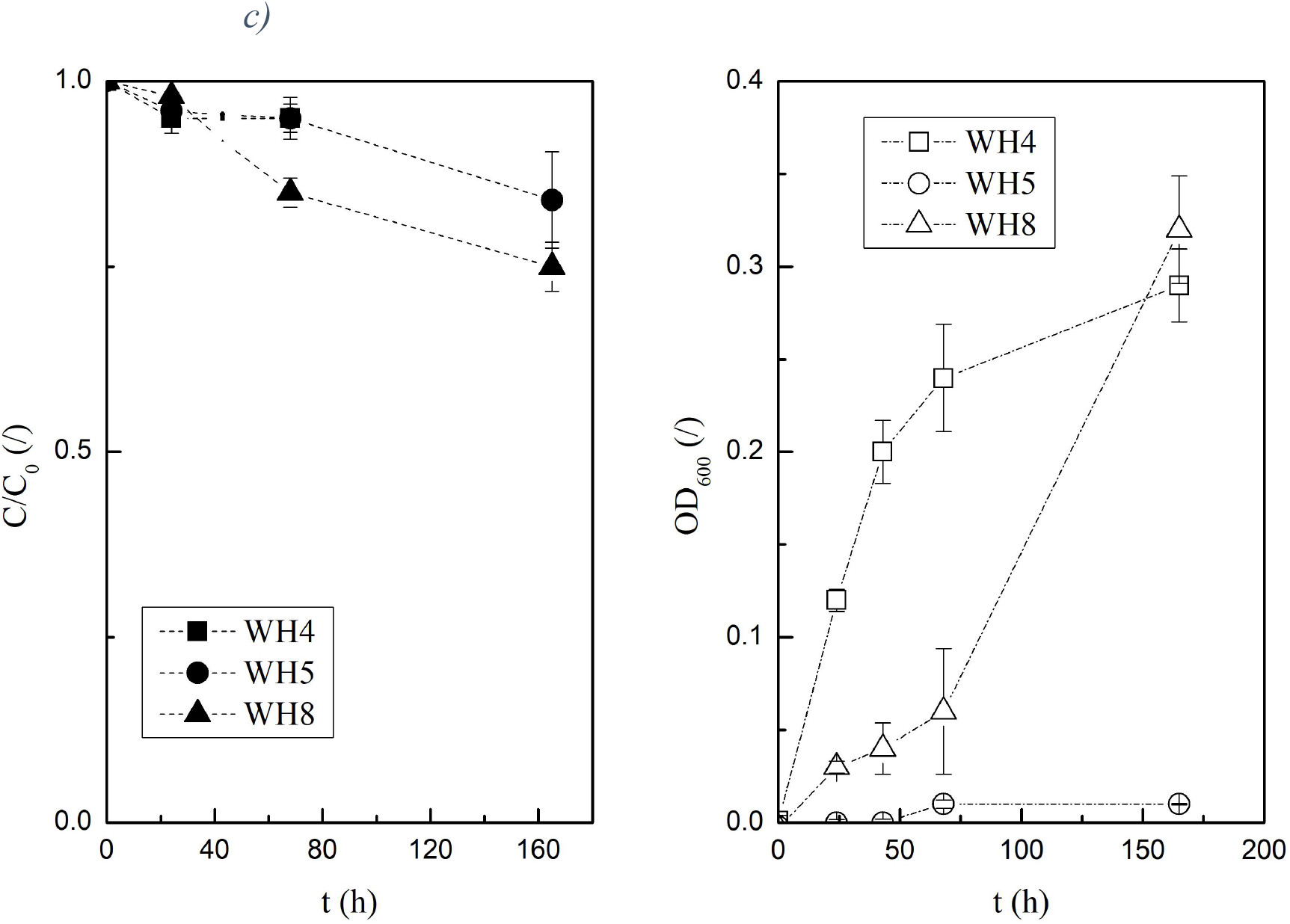
Degradation activity of the most active isolates for red, blue and whitener incubated up to 165 h at 25°C. Error bars indicate SE of triplicates; symbols C/C_0_ full and OD_600_ empty. a) BLU19, BLU19A and BLU23 in NB+10 mg/L blue; low values of C/C_0_ indicate high degradation activity. b) RED14, RED16B, RED14A and RED15A in NB+4.7 mg/L red; c) WH4, WH5 and WH8 in M9 glucose+0.5 g/L whitener.

As a result, we selected 16 of the best degrading strains for five dyes. Accordingly, in 24 h 2 strains degrading blue dye, BLU19 and BLU23, showed significant differences, degrading 40±3% and 33±2% C/C_0_ respectively (Fig. 3a). After 48 h, however, a linear decrease to 13%±0% C/C_0_ was achieved and no differences in degradation were observed between all strains. The discoloration of the red dye, which was 9-16 times faster than by other strains, was observed for strain RED14, which degraded 96% of the dye after 24 h (Fig 3.b). After 48 h there was no significant difference between the strains RED (14, 14A, 15A) that digested all the dye. Since we used a ~100-fold higher concentration of the whitener, this resulted in higher C/C_0_ values (Fig. 3c). Therefore, no direct comparison of the C/C_0_ values with degraders of red and blue was possible. Measurements of WH5 OD_600_ showed no growth in 165 h, which is in contrast to the cfu number and indicates an interference of the whitener with light absorption. The maximal degradation of whitener to 75% C/C_0_ in the first 72 h was recorded for WH8, which was 12% more than others. The isolates, degrading a yellow dye, were selected under two different protocols. The most active ones, Y17, Y18, Y19, degraded 72±3% C/C_0_ of the yellow dye at 72 h, but no significant differences between the strains were observed (Suppl. A, Table A.4, Fig. A.1d). Black dye was discolored only up to 88±3% average C/C_0_ by 3 isolates with OD_600_ 1.8-2.5 in 72 h. Since the degradation was too low, the time-dependent activities of the most active black isolates were not determined.

### 3.4 Principal component analysis of carbon source usage repertoire for the 62 isolates

By means of principal component analysis (PCA) we found out that the degraded compounds can be divided into three groups: whitener, the least biodegradable compound, azo dyes and readily biodegradable compounds (Fig. 4). No influence of the inoculation site on the degradability of the isolates was observed. Nevertheless, out of 62 selected isolates tested for repertoire of carbon source use 39%±9% from CF or BF and 73% of BTP-derived isolates are extreme degraders of at least one type of compound. From CF extreme degraders of readily biodegradable compounds and extreme degraders of dyes in glucose were found. From BF originated extreme degraders of whitener (25% of 20). From BTP we isolated *Aeromonas* sp. RES19, the best generalist, with an average of C_U_/C_0_ 80%±6% for all compounds. Extreme degraders have obtained above-average relative value of C_U_/C_0_>50% and for whitener >70% (Suppl. B, Table B.5).

**Fig. 4.**
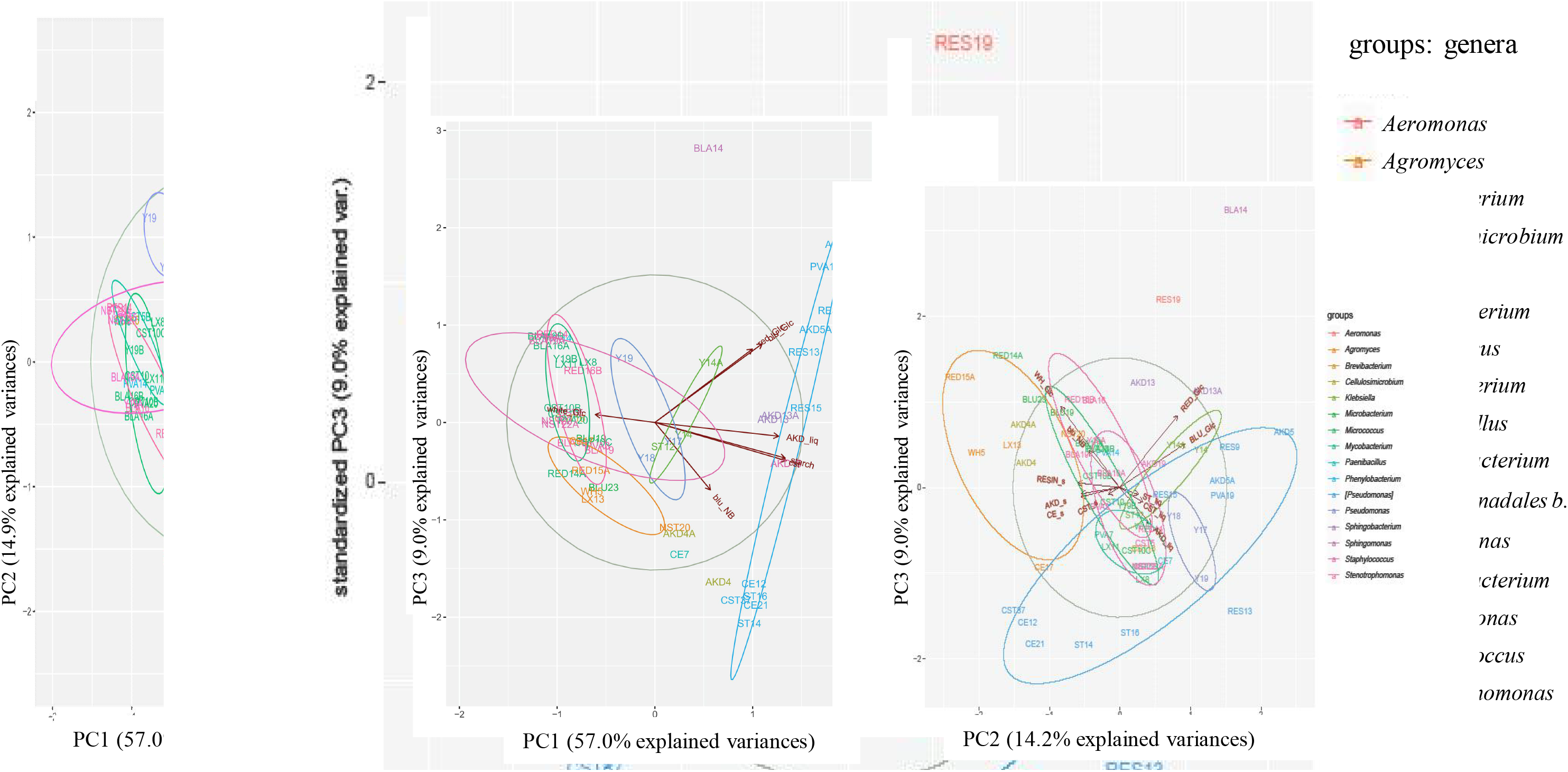
Results of the principal component analysis of the repertoire of carbon source consumption for 58 isolates in liquid media with clusters of the genera PC1PC2 (left) and PC1PC3 (right).

With 66% explained variances (PC1PC3 in Suppl. A, Fig. A.4), a positive correlation was found between the test results for CST or AKD in agar and liquid media with 76%±3% equal numbers of above- and below-average degraders. With 71.9 % explained variances (PC1PC2 in Suppl. A, Fig. A.4) we found a degradation activity of 34 isolates for cationic starch, AKD, resin and cellulose in agar media. 70% of the extreme cellulose degraders also degrade AKD, resin and CST, while with 60.5 % explained variances (Fig. 4) 68% of the starch degraders can also use AKD in liquid.

Isolates of the genus *Xanthomonadales bacterium* were characterised by the use of readily biodegradable compounds, with C_U_/C_0_ 81%±5%, but showed no activity for whitener. With 63% explained variances (PC1PC3 in Fig. 4) *Sphingomonas* sp. BLA14 was the highest strain for decolorization of dyes, with an average discoloration C_U_/C_0_ 80%±1%, but below-average decomposition of other compounds.

### 3.5 The 6 selected isolates

From 62 isolates we selected six (Fig. 5) that degrade starch, all readily biodegradable compounds, azo dyes in two different substrates, whitener and the most general degrader with the broadest repertoire of carbon sources. Based on the high hydrolysis of starch in liquid media (Fig. 5a) and whitener after 4 days with C_U_/C_0_ 100% and 72% respectively, the AKD4 strain was selected, with the closest type strains *Cellulosimicrobium funkei* (T) and *C. cellulans* (T) (Suppl. A, Fig. A.5c). *Xanthomonadales bacterium* strain CST37, with the closest type strain [*Pseudomonas*] *boreopolis* (T) (Suppl. A, Fig. A.5a), also hydrolyzed 100% starch. It was selected because it was most active for readily biodegradable compounds with an average of C_U_/C_0_ for starch, cellulose, resin acids and AKD 87%±16% (Fig. 5b). Due to the highest average activity for azo dyes in glucose (C_U_/C_0_ 55% blue and 100% red) BLA14 was selected, with the closest type strain *Sphingomonas paucimobilis* (T) (Suppl. A, Fig. A.5b). The strains Y14A and RED15A were most active (95%±3%) for blue dye in NB. Y14A, with the related strains *Klebsiella pneumoniae* (T) and *K. quasipneumoniae* (T) (Suppl. A, Fig. A.5d), could also use the yellow dye as the only carbon source. RED15A was selected since it degrades whitener above average (C_U_/C_0_ 86%) and is an average degrader of starch, resin acids, AKD and cellulose in agar media with average C_U_/C_0_ 64%±8%. It is sister strain of *Agromyces indicus* (T) (Suppl. A, Fig. A.5c). RES19, related to *Aeromonas dhakensis* (T) and *A. caviae* (T) (Fig. 6), was chosen as an extreme degrader of starch (C_U_/C_0_ 99%±1%) and degrader of whitener and red and blue dye with average C_U_/C_0_ 78%±5%.

**Fig. 5.**
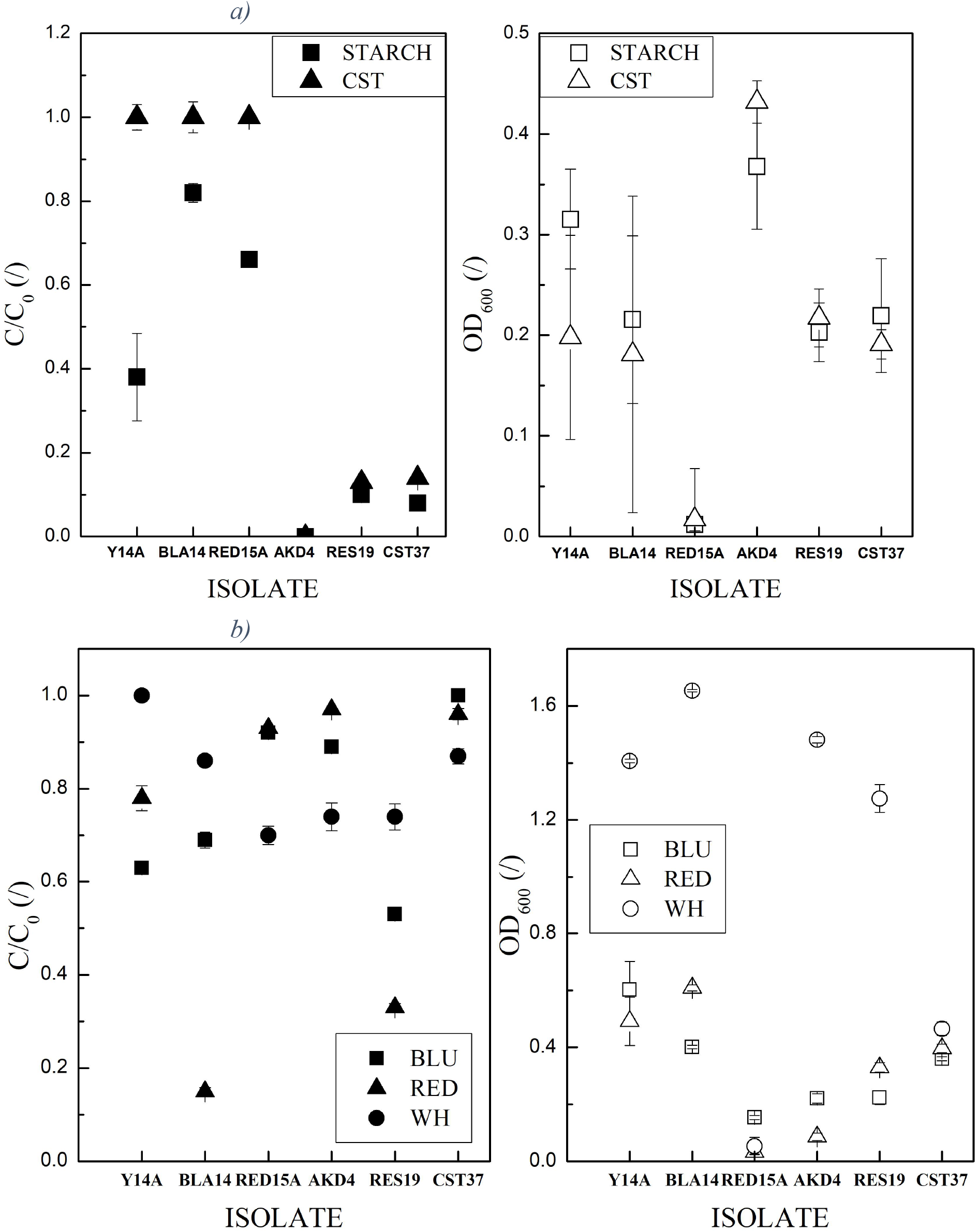
Reservoir of carbon sources of the six selected isolates. Error bars indicate RSE of triplicates. Low values of C/C_0_ indicate high degradation activity. C/C_0_ full and OD_600_ empty symbols. a) Starch and b) dyes and whitener in glucose.

**Fig. 6.**
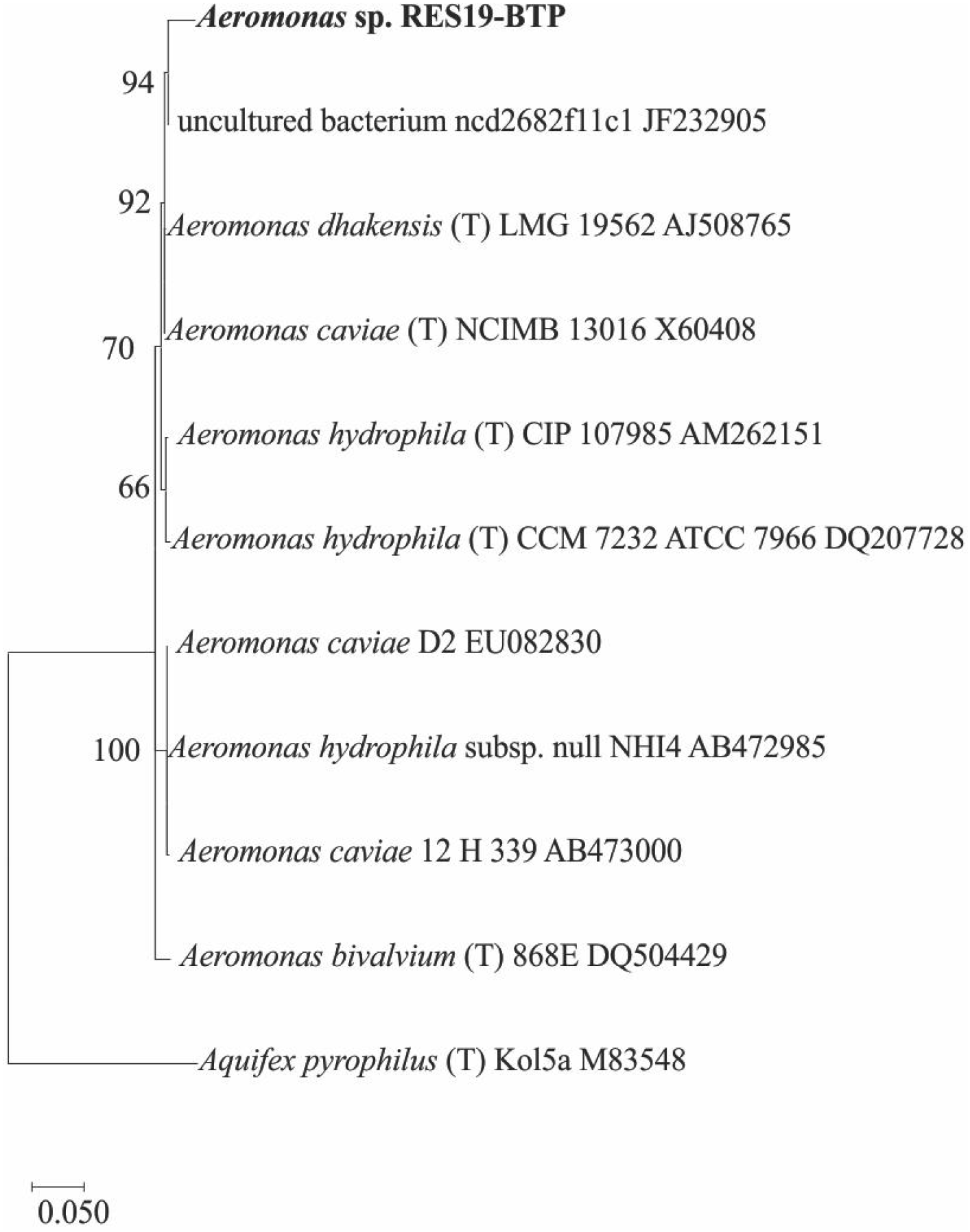
Phylogenetic tree displaying the phylogenetic position of RES19 and some related members of the genus Aeromonas. Aquifex pyrophilus (T) was used as an outlier. Numbers indicate percentages of bootstrap support, derived from 1000 resamplings.

### 3.6 The coexistence test

Coexistence in media with synthetic and real whitewater was compared for all 62 combinations of the 6 selected isolates (Suppl. B, Table B.6). After 3 days the individual isolates consumed less organic carbon than the combinations. In synthetic and real whitewater CST37 used 27% and 31% less than the best combination of 4, respectively. The combinations of 4 isolates were 23% more active in real whitewater than the single extreme general degrader RES19. Although the composition of synthetic whitewater was assumed from shares in the stock solution (Suppl. A, Table A.2), similar combinations of isolates were most active in both media. The most active combination in real media, with a C/C_0_ of 62.5±1.5% in both media, consisted of CST37, BLA14, RES19 and AKD4 (Fig. 5). Therefore, this consortium was considered the one with the highest potential for whitewater bioaugmentation. The relatively low levels of degradation were attributed to the minimal content of M9 salts in media with real and large amounts of the dye in synthetic whitewater.

### 3.7 Pilot test with an immobilized consortium of 4 isolates

To test the concept, a pilot test was conducted under non-sterile conditions. We used the best consortium of four isolates, CST37, BLA14, AKD4 and RES19, immobilized on the carriers, which were introduced into a 33-liter column (Suppl. A, Fig. A.6). After the flow had been stabilized, COD decreased from an initial 205 mg/L of inflow to 59 mg/L of outflow, corresponding to a 71% reduction of COD of filtered water. Initially, a large coefficient of 2.7 in COD between unfiltered and filtered effluent was found. As expected, after three days the difference became insignificant. On used carriers, taken 20 cm below the top of the column, 2.5×10^4^ cfu/mL RES19, CST37 and AKD4, no BLA14 and 11% unknown microorganisms were counted. On the 21^st^ day, COD of a new batch of whitewater decreased from 400 to 50 mg/L, i.e. by 88% (Suppl. A, Fig. A.6 and Suppl. B, Table B.7).

## 4 Discussion

We have fulfilled the aim of this study by confirming that the selected indigenous bacteria of the genera *Xanthomonadales bacterium*, *Sphingomonas*, *Cellulosimicrobium* and *Aeromonas* are capable of purifying the whitewater of a wood-free paper mill (Suppl. A, Fig. A.6). Whereas treatment by photolysis with UV irradiation and photocatalysis with titanium dioxide was ineffective (Suppl. A, Section A.1). Stable catabolism of the four selected isolates, probably developed under nutrient-poor conditions during isolation (Horemans *et al.*, 2017b), was confirmed with an 88% decrease in COD even on day 21 of the pilot experiment. The coexistence tests that gave the best combination of 4 bacteria (Suppl. B, Table B.6) confirmed results of (Santisi *et al.*, 2015) that more bacteria have an advantage over a single generalist. Already during the laboratory tests the four selected bacteria were among the most capable to decompose separate papermaking chemicals (Fig. 5). Two of the isolates are extreme degraders of different types of compounds (CST37 and BLA14), and two are general degraders (RES19 and AKD4).

The proof-of-concept and repertoire of carbon source use results in Suppl. B, Table B.5 and Table B.7 confirm our hypothesis that microbes from a wood-free paper mill are well suited for the use of the pollutants present. On the other hand, based on the selection results in Suppl. A, Fig. A.1 we can also confirm that “black box” microorganisms, taken from the same environment, would be a bad start-up inoculum for a biological treatment plant. Namely, the 355 isolates adhere to the Pareto principle (Dejonghe *et al.*, 2001). In most cases they are below average degraders of papermaking compounds.

Slow growth of bacterial isolates on isolation media with minimal amounts of papermaking compounds was confirming findings (Gray *et al.*, 2019) that extremely slow growth is an alternative strategy for bacteria to survive deep hunger. In addition, we detected an anomaly in the phylogenetic tree of identified isolates. Namely, the position of the 16 strains with the nearest relative, [*Pseudomonas*] *boreopolis* (T), is not as we have expected from its classification in Bac Dive (Fig. 2). However, it is consistent with the publication in IJSEM (Anzai *et al.*, 2000). The 16 strains have therefore been placed in the order Xanthomonadales as a new genus, the *Xanthomonadales bacterium*.

As we suspected, we were able to isolate degraders of readily- and non-readily biodegradable compounds. Accordingly, PCA results in Fig. 4 show that *Xanthomonadales bacterium* strains stand-out as highly active degraders of readily biodegradable papermaking compounds. On the other hand, RED14A is, in accordance with (Cui *et al.*, 2012), specialised in the recalcitrant whitener (Suppl. B, Table B.5).

The results for *Xanthomonadales bacterium* sp. strain CST37 were confirmed by the reported high activity of [*Pseudomonas*] *boreopolis* for PVA utilization and discoloration of dyes (Guo *et al.*, 2018). However, there are no previous reports on its utilization of AKD, resin acids and cellulose. The discoloration of dyes with *Klebsiella pneumoniae* (Zablocka-Godlewska *et al.*, 2015) and the use of polycyclic aromatic hydrocarbons by *Sphingomonas paucimobilis* (San Miguel *et al.*, 2009) are consistent with our results for *Klebsiella* sp. strain Y14A and *Sphingomonas* sp. strain BLA14. Lignocellulolytic enzymes secreted by *Cellulosimicrobium cellulans* (Liu *et al.*, 2015), starch hydrolysis by *Agromyces* sp. (Li *et al.*, 2003) and broad metabolic abilities of *Aeromonas hydrophila* (Seshadri *et al.*, 2006) are in accordance with our findings for *Cellulosimicrobium* sp. strain AKD4, *Agromyces* sp. strain RED15A and *Aeromonas* sp. strain RES19. Hence, reviewing the literature of the closest relatives of the 6 isolates selected by PCA confirmed accuracy of our selection methods (Suppl. A, Fig. A.2).

In addition, with PCA we found, in accordance with (Solís *et al.*, 2012), that the discoloration of the dyes depends on the presence of substrates. Namely, dye-degrading enzymes could be excreted stochastically in NB, while discoloration in glucose is more dye-specific. In addition, similar biodegradation activities found in selection results for starch or AKD in liquid and solid media (Suppl. A, Fig. A.4) indicate that inclusion of enzymes in 1.5% agar is not a rate limiting factor.

In short, the strategy for preparation of an artificial consortium of indigenous bacteria is discussed in the following section.

### 4.1 The strategy for preparation of consortium for bioaugmentation

1. take an environmental sample and isolate 300-400 bacteria (8 months); The isolation of the bacteria at low nutrient levels, comparable to 500-fold diluted lysogeny broth, was the most time-consuming step. Therefore, we propose a confirmatory study on the stability of catabolism in bacterial isolates in terms of accessibility of nutrients.
2. test the activity of isolates to degrade the targeted organic compounds (3 months); We disregarded any ambiguous selection methods and eliminated bacteria, like RED16B that in accordance with (Assih *et al.*, 2002) produces stained products in NB (Fig. 3), which might interfere with the treatment process. Since unambiguous selection methods for complex media are lacking, we propose to develop new analytical methods using specialist bacteria.
3. find the combinations that can best be implemented in the desired environment (1 month);
4. upscale the best combination in the desired environment for the proof of concept (3 months);
5. upscale for the industrial application (1 year); Based on the proof-of-concept results in Suppl. A, Fig. A.6 we assume that the transfer to the industrial environment would have a similarly high efficiency. However, before final application we expect at least one more year of optimisation of bioprocess reactor design in combination with 30-day pilot experiments (Tiirola *et al.*, 2003).

## 5 Conclusions

355 isolated bacteria adapted to the wood-free paper production process were mostly below average degraders of the tested organic compounds. They were isolated from three interconnected inoculum sources: whitewater, biofilm from the paper machine and wastewater from the biological treatment plant. In most cases, the inoculum sources determined the genera of the isolated bacteria.

By principal component analysis, the degradation activities for papermaking chemicals of 58 isolates were investigated with respect to the genera. We found the following new findings: (a) In most cases, the starch degraders are also active in the degradation of alkyl ketene dimers. (b) The discoloration of dyes in different substrates is uncorrelated. (c) Whitening agents are degraded by a different mechanism than azo dyes and readily biodegradable compounds. (d) *Xanthomonadales bacterium* sp. is characterized by a high activity for readily biodegradable compounds.

The four bacteria selected for bioaugmentation are two specialists, *Xanthomonadales bacterium* and *Sphingomonas* sp., which complement each other in various niches, and two general degraders, *Cellulosimicrobium* and *Aeromonas* sp.

To prove the concept, the four highly active bacteria were immobilized on carriers and introduced into a 33-L column filled with whitewater. On day 21 an 88% decrease of COD was measured at a retention time of about 15 h. Based on the results obtained, these bacteria are able to purify whitewater of a wood-free paper mill, in contrast to treatment by photolysis with UV irradiation and photocatalysis with titanium dioxide, which was ineffective.

The application of such a bioaugmentation strategy would not only significantly reduce water consumption during the paper manufacturing process itself, but also increase productivity and reduce the amount of organic compounds released into the environment.

## Abbreviations

16S rRNA: 16S ribosomal ribonucleic acid
cfu: colony forming units
COD:N:P: chemical oxygen demand: total nitrogen: total phosphorus
DOC: dissolved organic carbon
OD_600_: absorption at 600 nm
PCA: principal component analysis
PCR: *Polymerase* chain reaction
PVA: polyvinyl alcohol
RDP: Ribosomal Database Project
SE: standard error

## Notation

A_CELLS_: absorption of bacteria
AKD: alkyl ketene dimer
BF: biofilm
BLA: black dye HM 2482
BLU: blue dye
BTP: effluent from the biological treatment plant
C/C_0_: coefficient between concentration of unutilized vs. initial carbon source
C_U_/C_0_: coefficient between utilized vs. initial carbon source in the relative range 0-1
CE: cellulose
CF: clear filtrate
CST: cationic starch
DB15: direct blue 15 dye
Glc: glucose
LX: latex
NB: nutrient broth No.1
NST: native starch
P&P: pulp and paper
RED: direct red 253 dye
RES: resin acids
ST: starch
WH: whitener
Y: yellow dye

## Acknowledgements

The authors acknowledge financial support from the Ministry of Education, Science and Sport of the Republic of Slovenia (research programme No. P2-0150) and European Regional Development Fund (ERDF) within the Operational Program for the Implementation of European Cohesion Policy in the period 2014–2020, Slovenian national projects (J4-7640, J1-6746, J3-1762, J1-9194, J7-9400 and P1-0143), Flemish-Slovenian research project: Bioavailable mercury methylation in the Adriatic sea (BE MERMAiD, grant agreement N1-0100), project CROSSING (grant PIE-0007,) European Urban Initiative Actions founded project Applause (Grant agreement UIA02-228), 2019 - 2023 (EU - Horizon 2020): InteGRated systems for Effective ENvironmEntal Remediation (GREENER, grant agreement 826312) and European Commission (SurfBio project, grant no.: 952379). The authors thank J. Jelnikar and Š. Božič for their support and J. Teržan and P. Djinović for proofreading and comments.

