## Supplemental Material A for "Isolation, identification and selection of bacteria with the proof-of-concept for bioaugmentation of whitewater from woodfree paper mills"

### SUPPLEMENTARY INFORMATION A

#### A.1 Photolytic and photocatalytic experiments

The photolytic and photocatalytic degradation of whitewater was investigated in a 120 mL batch reactor at atmospheric pressure. The whitewater was filtered through a membrane filter (Sartorius, 0.45 μm) before use. The reactor was controlled by a thermostat at 40°C (Julabo, model F25/ME) and magnetically stirred (300 rpm). The photocatalytic oxidation run was performed with a standard titanium dioxide photocatalyst, Degussa P25 (Evonik). P25 is widely used in many photocatalytic reactions due to its relatively high activity (Ohtani *et al.*, 2010). The catalyst concentration was in the range of 0-200 mg/L (photolytic or photocatalytic experiments). Before the illumination time, the suspension with the catalyst was kept in the dark (30 min) to allow the sorptium equilibrium to be established. The reactor contents were illuminated with a UV-C lamp (max. at $\lambda$ = 254 nm). The lamp was positioned in a quartz mantle, which was placed vertically in the middle of the reactor. Representative 2 mL samples were periodically taken from the reactor suspension for subsequent analysis of the total organic carbon (TOC) content. The samples were filtered through a membrane filter to remove catalyst particles. A CHNS analyzer (Perkin Elmer, model 2400 Series II) was used to determine the carbon content accumulated on the surface of the used catalyst.

Although the process water was filtered through a membrane filter (0.45 μm) before the experiments, the actual TOC difference after two hours of photocatalytic oxidation run was close to 0%. In fact, the reduction in TOC was ~ equal to the amount of adsorbed carbon on the catalyst surface obtained by CHNS analysis of the used catalyst. Therefore, we concluded that the suspended organic matter strongly adsorbed on the catalyst surface and blocked active sites. When the pH value of the process water was lowered to 1.5 by $H_2SO_4$, an 89% difference in TOC was measured after two hours of irradiation. Nevertheless, these results are due to the necessary condition to obtain constant pH value unsuitable for treatment of large amounts of whitewater.

### A.2 Methods development for selection of bacteria

#### A.2.1 Readily biodegradable compounds

From the discoloration radius ($r_{DIS}$), the concentration of the unused organic compound ($C$) and the media layer thickness (δ), the approximate mass of the C source used (mU) can be calculated using the equation $m_U = \pi\ r_{DIS}^2\ \delta\ C_C$. Therefore, the $C/C_0$ values were calculated using the following equation (Eq. A.1):

$$C/C_0=(r_{DIS*}^2- r_{DIS}^2 )/r_{DIS*}^2$$

$r_{DIS}$* is the discoloration radius of the best degrader strain with the total discolored content.

#### A.2.2 Azo dyes and whitener

First, influence of centrifugation on sedimentation of dyes was tested by determining the absorption or emission intensity of the dyes before and after centrifugation. For whitener, excitation at 315 nm showed a maximum wavelength emission at 440 nm, as for 4,4'-diamino-2,2'-stilbenedisulfonic acid (DSDS), and the coefficient between the fluorescence intensity before and after centrifugation was 97%. Black dye showed an absorption decrease of 4% at both peak maxima, red, blue and direct blue 15 (DB15) no concentration decrease and yellow dye showed a decrease of 50%. For dyes and whitener no concentration dependence of the maximum peak position was found. Calibration curves for all tests in liquid media were generated for molar concentrations at $\lambda_{max}$, and the volume of liquid in microplates was kept constant to ensure accurate absorbance measurements. To quantitatively determine the amount of unhydrolyzed dye by measuring the absorption of supernatants, the tests with NB in microplates had to be completed in a maximum of three days. To determine the activities of the most active bacteria, initially, relative standard error between the triplicates for all isolates in NB + dye and M9 glucose (Glc) + whitener was maximum 3%; M9 Glc means M9 mineral media supplemented with 4 g/L glucose. The microtiter plates were covered with sterile lids that were packed in bags to prevent evaporation. Wet paper was added to the bags to maintain a constant humidity. They were shaken gently at less than 50 rpm.

Two protocols were used for the determination of the non-degraded yellow dye, as cell removal could not be performed before determining the amount of non-degraded dye. For the first protocol the colonies were transferred in triplicates to NB, the four fastest growing cells were purified (three times at 5000 rpm in 9 g/L NaCl) and incubated in M9 yellow (10 μL of purified cells in NaCl transferred to 200 μL M9 yellow). After 48 hours the absorbance at 400 nm was recorded and the CFU were counted. For the second protocol, a bacterial colony was picked and transferred to mineral media supplemented with 82 mg/L yellow or 47 mg/L glucose. Incubation in mineral media with 47 mg/L glucose was used to reflect the absorption of the cells. After 168 hours of gentle shaking at less than 50 rpm and room temperature, the absorbance was recorded at 400 and 600 nm.

#### A.3 Selection of bacteria in media with dye as sole carbon source

In media with 47-95 mg/L dye as the only carbon source, no visual discoloration and no increase in $OD_{600}$ was observed in 10-13 days. Due to the 5% decrease in intensity of sterile media with time and the only slight decrease in intensity in inoculated media, no clear results were obtained. For example, after 13 days in M9 BLU at 30 °C *Pseudomonas* sp. BLU24- BF, *Acinetobacter* sp. BLU27-BTP, *Rhabdobacter* sp. BLU8- BF and *Pseudomonas* sp. BLU15- CF $C/C_0$_560 was 80%±1%, 87%±3%, 88%±1% and 89%±0%, respectively (calibration curve $C=0.0486\times A_{560}-8.28\times10^{-4}$). After 11 days in M9 BLA only *Stenotrophomonas* sp. BLA19- BF discolored black to 82%±5% $C/C_0$_590.

*Table A.1*

*Whitewater analysis of the wood-free paper mill within a period of two months with average, minimum and maximum values.*

| **N** | **COD (mg/L)** | **DOC (mg/L)** | **$BOD_5$ (mg/L)** | **$HCO_3^-$ (mg/L)** | **$Cl^-$ (mg/L)** | **$NO_3^-$ (mg/L)** | **$SO_4^{2-}$ (mg/L)** | **$Na^+$ (mg/L)** | **$K^+$ (mg/L)** | **$Ca^{2+}$ (mg/L)** | **$Mg^{2+}$ (mg/L)** | **pH** |
|---|---|---|---|---|---|---|---|---|---|---|---|---|
| **AVG** | **303** | **94** | **166** | **153** | **40** | **8** | **179** | **78** | **0** | **59** | **15** | **7.5** |
| **MIN** | **190** | **31** | **80** | **50** | **25** | **1.6** | **109** | **62** | **0** | **41** | **10** | **6.8** |
| **MAX** | **500** | **175** | **254** | **269** | **62** | **15** | **309** | **95** | **1.5** | **88** | **29** | **8.2** |
| **-1** | 232 | 134 | 158 | 165 | 49 | 7.8 | 183 | 78 | 0 | 52 | 16 | 7 |
| **1** | 214 | 54 | | 130 | 34 | 15 | 175 | 62 | 0 | 54 | 16 | 7.1 |
| **2** | 278 | 52 | 80 | 114 | 25 | 13 | 309 | 67 | 0 | 88 | 17 | 7.5 |
| **5** | 190 | 36 | | 111 | 26 | 7.4 | 267 | 90 | 0 | 78 | 13 | 7.2 |
| **6** | 210 | 31 | | 90 | 26 | 8.4 | 274 | 86 | 0 | 79 | 16 | 7.5 |
| **7** | 200 | 37 | | 50 | 26 | 9.1 | 233 | 70 | 0 | 59 | 14 | 6.8 |
| **8** | 268 | 77 | 109 | 141 | 36 | 9.1 | 233 | 95 | 0 | 67 | 14 | 7.5 |
| **9** | 320 | 91 | | 156 | 45 | 3.7 | 132 | 66 | 0 | 43 | 17 | 7.3 |
| **12** | 252 | 78 | | 123 | 39 | 5.7 | 117 | 66 | 0 | 45 | 12 | 7.6 |
| **13** | 356 | 95 | | 144 | 36 | 7.7 | 156 | 78 | 0 | 52 | 13 | 7.6 |
| **14** | 300 | 120 | 178 | 184 | 46 | 8.2 | 125 | 85 | 0 | 51 | 13 | 7.4 |
| **15** | 284 | 105 | | 180 | 41 | 4 | 138 | 76 | 0 | 64 | 14 | 7.8 |
| **16** | 345 | 114 | | 170 | 44 | 6 | 109 | 75 | 0 | 41 | 10 | 7.7 |
| **21** | 362 | 128 | | 141 | 36 | 6.2 | 150 | 76 | 0 | 60 | 13 | 7.5 |
| **22** | 500 | 175 | 180 | 168 | 44 | 2.3 | 116 | 76 | 0 | 50 | 11 | 7.3 |
| **24** | 243 | 74 | | 167 | 27 | 4.3 | 122 | 76 | 0 | 60 | 12 | 7.4 |
| **28** | 448 | 128 | 183 | 146 | 57 | 13 | 235 | 84 | 0 | 61 | 16 | 8.2 |
| **31** | 393 | 120 | 254 | 253 | 52 | 11.1 | 171 | 82 | 0 | 52 | 14 | 7.4 |

| | | | | | | | | | | | | |
|---|---|---|---|---|---|---|---|---|---|---|---|---|
| **59** | 371 | 145 | 183 | 269 | 62 | 1.6 | 163 | 87 | 1.5 | 71 | 29 | 8.2 |

*Table A.2*

*Organic substances used as carbon sources with abbreviation, approximate proportions in the stock solution and chemical structure with molar mass to calculate the calibration curves; for dyes maximum wavelength absorption or excitation/emission and dye intensity before and after centrifugation at 12,000g.*

| Abbr. | Share (%) | Chemical structure |
|---|---|---|
| CST | 0.4 | Cationic potato starch, degree of derivatisation 0.04, based on integrals in $^{1}$H-NMR; M($C_6H_{10}O_5$)=162 g/mol (monomeric unit) |
| NST | <0.001 | Native starch, CAS 9005-25-8; (2R,SS,4S,5R,6R)-2-(hydroxy methyl)-6-[(2R(xy-oxane-3,4,5-triol,3S,4R,5R,6S)-4,5,6-tri hydroxy-2-(hydroxymethyl)oxan-3-y]oxy-oxane-3,4,5-triol; M($C_6H_{10}O_5$)=162 g/mol (monomeric unit) |
| ST | <0.001 | Starch, soluble (Sigma Aldrich, USA) CAS 9005-84-9; M($C_6H_{10}O_5$)=162 g/mol (monomeric unit) |
| CE | 4 | Microcrystalline cellulose (VWR, USA) CAS 9004-34-6; M ($C_{12}H_{22}O_{11}$)=342 g/mol (monomeric unit) |
| PVA | <0.001 | Polyvinyl alcohol, CAS 9002-89-5; M($C_2H_4O$)=44 g/mol (monomeric unit) |
| LX | <0.001 | Latex dispersion, styrene-butadiene copolymer; M($C_{12}H_{14}$)=158 g/mol (monomeric unit) |
| RES | 0.4 | Rosin, CAS No 8050-09-7; contains abietic and pimeric type resin acids; M($C_{20}H_{30}O_2$)=303 g/mol |
| AKD | 0.18 | Alkyl ketene dimers: 2-oxetanone, 3-C12-16-alkyl-4-C13-17-alkylidene derivatives, CAS No 84989-41-3; M=477 g/mol |
| WH | 0.10 | Whitening agent; fluorescent brightening agent; tetrasodium 2,2'-ethene-1,2-diylbis[5-({4-[bis(2-hydroxyethyl)amino] -6-[(4-sulfonatophenyl)amino]-1,3,5-triazin-2-yl}amino]benzenesulfonate]; spectral properties similar to 4,4'-diamino-2,2'-stilbenedisulfonic acid; M($C_{42}H_{44}N_{12}Na_4O_{16}S_4$)=1193 g/mol; excitation at 315 nm with maximum emission at 440 nm; 97% $I_{cent}/I$ |

| | | |
|---|---|---|
| BLA | 0.003 | Black dye HM 2482, CAS No. 89857-06-7; 1,3'-bipyridinium,5'-[[4-[[7-[(1',2'-dihydro-6'-hydroxy-3,4-dimethyl-2'-oxo[1,3'-bipyridinium]-5'-yl)azo]-1-hydroxy-3-sulpho-2-naphthalenyl]azo]phenyl]azo]-1'-[3-(dimethylamino)propyl]-1',2'-oxo-, salt with 2-hydroxypropanoic acid; M($C_{50}H_{57}N_{11}O_{14}S_4$)=1068 g/mol; $\lambda_{max}$=440, 590 nm; 96% $I_{cent}/I$ |
| Y | 0.003 | Yellow dye containing C; H; N; O; S; stilbene derivative; M [a]=180 g/mol; $\lambda_{max}$=400 nm; 50% $I_{cent}/I$ |
| RED | 0.003 | Direct red dye 253 (Marshall and Albert 1992), CAS 142985-51-1: Dinatrium;(3*E*)-7-[[4,6-bis(2-hydroxyethylamino)-1,3,5-triazin-2-yl]amino]-4-oxo-3-[[4-[(4-sulfonatophenyl)diazenyl]phenyl] hydrazinylidene] naphthalene-2-sulfonate; M($C_{29}H_{26}N_{10}Na_2O_9S_2$)=769 g/mol; $\lambda_{max}$=510 nm |
| BLU | 0.003 | Blue dye, similar biodegradation to direct blue 15, CAS No. 2429-74-5. To evaluate the structural similarity of the dye BLU to DB15, cometabolism tests on 44 selected strains were compared for the biodegradation of BLU and DB15 dye in NB. After 72 h similar discoloration patterns were found. Direct blue 15 is 2,3,3'-[(3,3'-dimethoxy-4,4'-biphenylylene)bis(azo)]bis [5-amino-4-hydroxy-tetrasodium salt]; $M_{DB15}$($C_{34}H_{28}N_6NaO_{16}S_4$)=927 g/mol; $\lambda_{max\ BLU}$=560 nm; $\lambda_{max\ DB15}$=600 nm |

[a] Unknown structure, molar weight taken for glucose.

*Table A.3*

*Protocols for the selection of bacteria for construction of consortium with high potential for bioaugmentation of whitewater with inoculum preparation and composition of media.*

| **Test** | **Inoculum preparation** | **Carbon source** | **Selection media** |
|---|---|---|---|
| Selection of isolates for the usage of carbon source which the particular isolate utilized during isolation | Colony of the isolate on NB agar was picked and transferred to a circle on selection media. Incubation was at 25 °C for 7-8 days. | starch | M9 10 g/L ST |
| | | | M9 10 g/L CST |
| | | | M9 10 g/L NST |
| | | CE | M9 5 g/L CE |
| | | RES | M9 0.25 g/L RES |
| | | AKD | M9 0.25g/L AKD |
| | | PVA | M9 5g/L PVA |
| | | LX | M9 0.25g/L LX |
| | Colony of the isolate on NB agar was picked and transferred to appropriate M9 | BLU | M9 + 47 mg/L BLU<br>NB + 5.7 mg/L BLU |

| | | | |
|---|---|---|---|
| | media and incubated at 25 °C for 3 days. 10 μL of the solution were transferred into the selection medium. | RED | M9 47 mg/L RED<br>NB + 15 mg/L RED |
| | | BLA | M9 95 mg/L BLA<br>NB + 57 mg/L BLA |
| | | WH | M9 0.9 g/L WH+4 g/L glucose |
| | | Y | first in NB, than in M9 37 mg/L Y (protocol 1) |
| | | | M9 82 mg/L Y / M9 47 mg/L Glc (protocol 2) |
| | | | NB+ 7.5 mg/L Y (cometabolism) |
| Time-dependent activity determination | A colony was transferred into corresponding M9 dye at 47 mg/L for 3 days at 25 °C. | RED | NB + 13 mg/L RED |
| | | BLU | NB + 10 mg/L BLU |
| | | WH | M9 4 g/L Glc + 0.5 g/L WH |
| Tests for the repertoire of carbon source usage | A colony was transferred into M9 Glc for 3 days at 25 °C. | CST | M9 0.5 g/L CST |
| | | ST | M9 0.5 g/L ST |
| | | AKD | M9 4 g/L AKD |
| | | RED | M9 Glc + 4.7 mg/L RED |
| | | BLU | M9 Glc + 4.7 mg/L BLU |
| | | WH | M9 Glc + 4.7 mg/L WH |
| | | BLU | NB + 4.7 mg/L BLU |
| | | CE | 20 mL / plate M9 5 g/L CE |
| | | CST | 15 mL / plate M9 5 g/L CST |
| | | AKD | 20 mL / plate M9 0.25 g/L AKD |
| | | RES | 20 mL / plate M9 0.25 g/L RES |
| Coexistence test | The six selected isolates were calibrated for cfu vs. $OD_{600}$. Overnight culture was harvested in NB, cleared three times in 9 | CST, RES, PVA, DB15 | M9 (CST, PVA, RES, DB15); in the ratio CST : PVA : RES : DB15 = 1 : 1 : 1 : 0.5 |

| | | | |
|---|---|---|---|
| | g/L NaCl. $OD_{600}$ was measured and a calibration curve was generated. The amount of volume was adjusted so that an equal initial sum of bacteria per each medium was added, i.e. $10^7$ cfu/mL. | whitewater, sterile filtered through 0.2 µm PP filter (Chromafil) | M9 sterile whitewater |
| Pilot test | Separate overnight cultures of the four bacteria of the consortium harvested in NB, cleared in 9 g/L NaCl, $OD_{600}$ measured and cfu calculated. | whitewater | whitewater (as is) with added urea and $H_3PO_4$ in the ratio COD : N : P = 100 : 5 : 1 |

*Table A.4*

*Selected isolates, active for the carbon source which they used during isolation with number of isolated bacteria per carbon source, $C/C_0$ and $OD_{600}$ values and biodegradation assay duration.*

| Carbon source | $N_0$ | Active isolate | $C/C_0$ (%)±RSE(%) | $OD_{600}$ (/)±RSE(%) | t (d) | $C_0$ (g/L) |
|---|---|---|---|---|---|---|
| CST | 50 | CST37 | 75 | / | 9 | 10 |
| | | CST5 | 83 | / | 15 | |
| | | CST10 | 83 | / | | |
| | | CST16 | 83 | / | | |
| NST | 32 | NST20 | 83 | / | 11 | 10 |
| | | NST22 | 87 | / | | |
| ST | 36 | ST12 | 72 | / | 13 | 10 |
| | | ST9 | 56 | / | | |
| | | ST14 | 0 | / | | |
| | | ST16 | 0 | / | | |
| RES | 20 | RES19 | 39 | / | 7 | 0.25 |
| | | RES9 | 75 | / | | |
| | | RES13 | 75 | / | | |
| | | RES15 | 75 | / | | |
| | | RES16 | 75 | / | | |
| AKD | 31 | AKD4 | 51 | / | 7 | 0.25 |
| | | AKD5 | 46 | / | | |
| | | AKD13 | 75 | / | | |
| | | AKD19 | 75 | / | | |
| PVA | 21 | PVA19 | 13 | / | 8 | 5 |
| | | PVA20 | 75 | / | 7 | |
| | | PVA3[a] | 75 | / | | |
| | | PVA7[a] | 75 | / | | |
| | | PVA14[a] | 84 | / | | |
| CE | 24 | CE12 | 0 | / | 11 | 5 |
| | | CE21 | 0 | / | | |
| | | CE7 | 81 | / | | |
| | | CE17 | 46 | / | | |
| LX | 23 | LX13 | 31 | / | 7 | 0.25 |
| | | LX11 | 75 | / | | |
| | | LX8 | 84 | / | | |
| RED | 24 | RED14 | -15±1.0 | 1.2± 0.5 | 3 | 0.013 |
| | | RED14A | -14±3.3 | 1.3± 8.5 | | |
| | | RED15A | -16±0.6 | 1.1±7.8 | | |
| | | RED16B | 68±5.5 | 1.8±5.4 | | |
| BLU | 27 | BLU19 | 13±1.5 | 0.9±0.5 | 3 | 0.01 |

| | | | | | | |
|---|---|---|---|---|---|---|
| | | BLU23 | 12±2.0 | 0.8±3.3 | | |
| WH | 13 | WH8 | 75±3.3 | 0.06±4.2 | 3 | 0.5 |
| | | WH5 | 84±6.5 | 0.01±2.3 | | |
| | | WH4 | / | 0.24±2.9 | | |
| Y | 26 | Y17 | 70±1 | 2.6E+08 [b] | 3[b] | 0.037 |
| | | Y14 | 62±18 | 4.9E+09 [b] | | |
| | | Y18 | 72±0.5 | 2.8E+13 [b] | | |
| | | Y19 | 75±3 | 4.8E+13 [b] | | |
| BLA | 33 | BLA14 | 70 | 1.8±2.1 | 2.8 | 0.057 |
| | | BLA16 | 70 | 2.4±2.7 | | |
| | | BLA19[c] | 81±3 | / | 11 | 0.095 |

[a] …no circled discoloring; [b]… due to the absorption of yellow dye particles the $OD_{600}$ values are unrealistic, therefore cfu are presented - protocol 1; [c]… incubation in M9 with black dye as the sole carbon source.

*Table A.5*

*Comparison of the absorption measurements with the results from DOC to confirm the accuracy of the AKD selection method; the bacteria were inoculated for 62 h, initially $10^8$ cfu/mL and $C_0$ 4 g/L.*

| **Isolate** | **$C/C_0$ (%) ± RSE (%)** | |
|---|---|---|
| | **UV-VIS spectrophotometry results** | **DOC results** |
| CST37 | 56±0 | 62±0 |
| AKD4 | 57±0 | 66±1 |
| RES19 | 56±0 | 71±1 |
| Y14A | 100±1 | 100±1 |

*Table A.6*

*Results of the time-dependent activity of the most active red, blue and fluorescent dye-whitener degraders for the dye they used during isolation at 25 °C in triplicates. The Cfu value of WH4, WH5 and WH8 after 165 h was $7\times10^8$, $10^9$ and $6\times10^{10}$ cfu/mL, respectively.*

| **Isolate** | **$C/C_{0\,\lambda max}$ (%)±RSE (%)** | | | **$OD_{600}$ (/)±RSE (%)** | | | | |
|---|---|---|---|---|---|---|---|---|
| | **24 h** | **48 h** | **72 h** | **0 h** | **24 h** | **48 h** | **72 h** | **165 h** |
| **BLU19** | 40±3.3 | 19±0.7 | 13±1.5 | 0.01±0.2 | 0.56±1.0 | 0.88±2.1 | 0.91±0.5 | / |
| **BLU23** | 33±2.1 | 17±1.4 | 12±2.0 | 0.01±0.4 | 0.60±0.7 | 0.88±3.0 | 0.82±3.3 | / |
| **RED14** | 6±2.5 | -7±2.2 | -15±1.0 | 0.01±0.6 | 0.1±0.9 | 1.3±3.6 | 1.2±0.5 | / |

| | | | | | | | | |
|---|---|---|---|---|---|---|---|---|
| **RED14A** | 103±0.4 | 2.8±3.1 | -14±3.3 | 0.02±1.8 | 0.1±6.4 | 1.1±1.5 | 1.3±8.5 | / |
| **RED15A** | 60±3.4 | -16±0.3 | -16±0.6 | 0.01±0.5 | 0.6±6.2 | 1.1±1.3 | 1.1±7.8 | / |
| **RED16B** | 65±4.0 | 35±1.6 | 68±5.5 | 0.01±1.0 | 0.1±6.5 | 1.6±5.4 | 1.8±5.4 | / |
| **WH8** | 98±3.1 | 85±2.0 | 75±3.3 | -0.003±1 | 0.03±2.0 | 0.04±10 | 0.06±4.2 | 0.32±10 |
| **WH5** | 96±1.9 | 95±1.9 | 84±6.5 | 0.00±0.4 | 0.00±1.8 | 0.00±2.1 | 0.01±2.3 | 0.01±2.7 |
| **WH4** | 95±2.0 | 95±2.2 | / | 0.00 | 0.18±0.6 | 0.19±1.7 | 0.24±2.9 | 0.29±5.1 |

a)

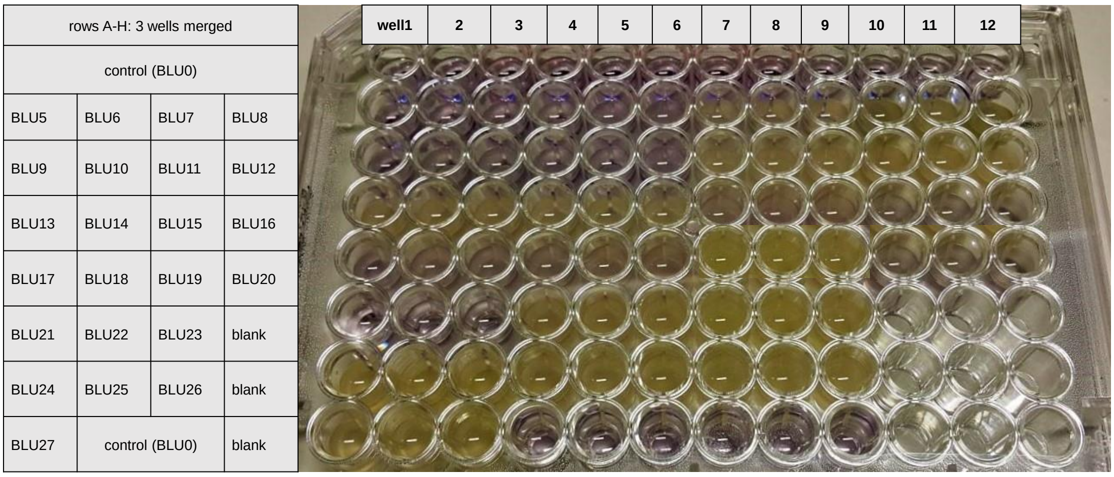

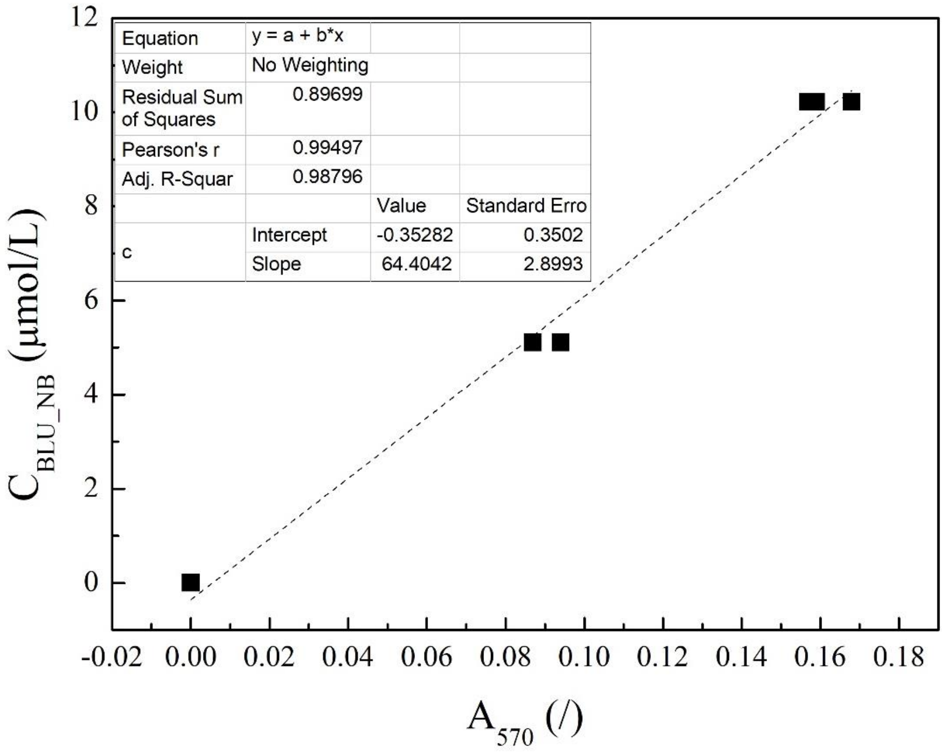
Equation
y = a + b*x
Weight
No Weighting
Residual Sum of Squares
0.89699
Pearson's r
0.99497
Adj. R-Squar
0.98796
Value
Standard Erro
c
Intercept
-0.35282
0.3502
Slope
64.4042
2.8993
$C_{BLU\_NB}$ (µmol/L)
$A_{570}$ (/)

b)

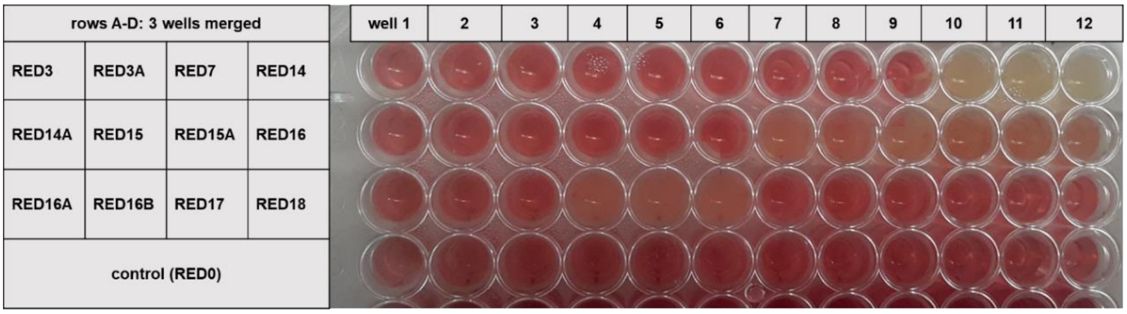
rows A-D: 3 wells merged
RED3
RED3A
RED7
RED14
RED14A
RED15
RED15A
RED16
RED16A
RED16B
RED17
RED18
control (RED0)
well 1

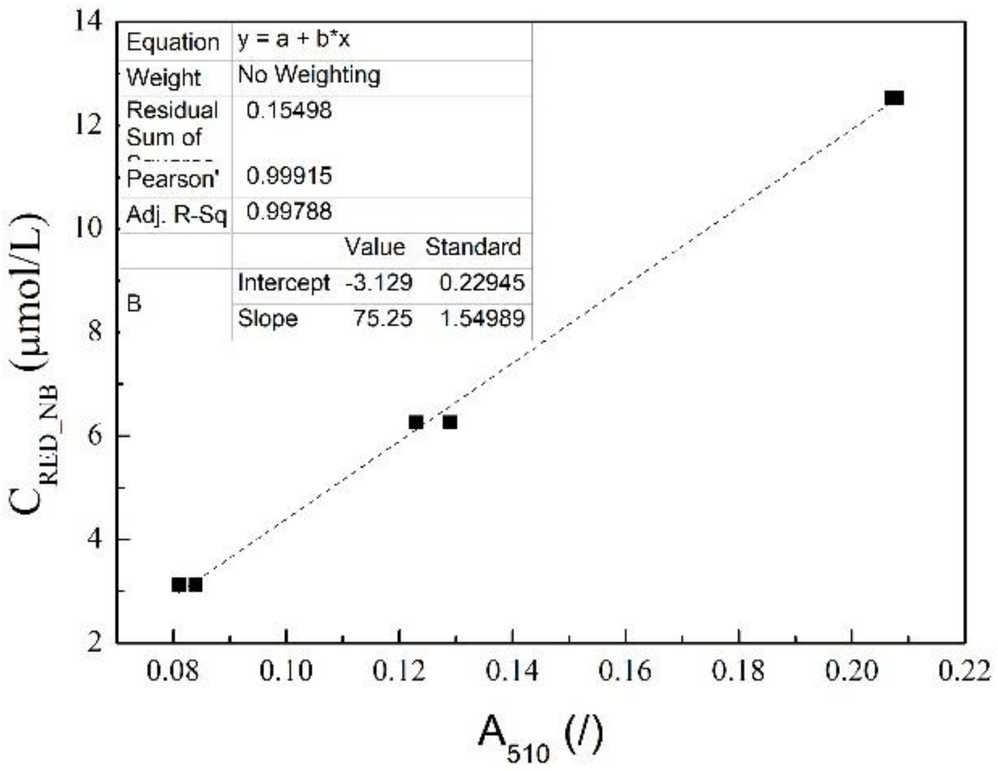
Equation
y = a + b*x
Weight
No Weighting
Residual Sum of
0.15498
Pearson'
0.99915
Adj. R-Sq
0.99788
Value
Standard
B
Intercept
-3.129
0.22945
Slope
75.25
1.54989
$C_{RED\_NB}$ (μmol/L)
$A_{510}$ (/)
14
12
10
8
6
4
2
0.08
0.10
0.12
0.14
0.16
0.18
0.20
0.22

c)

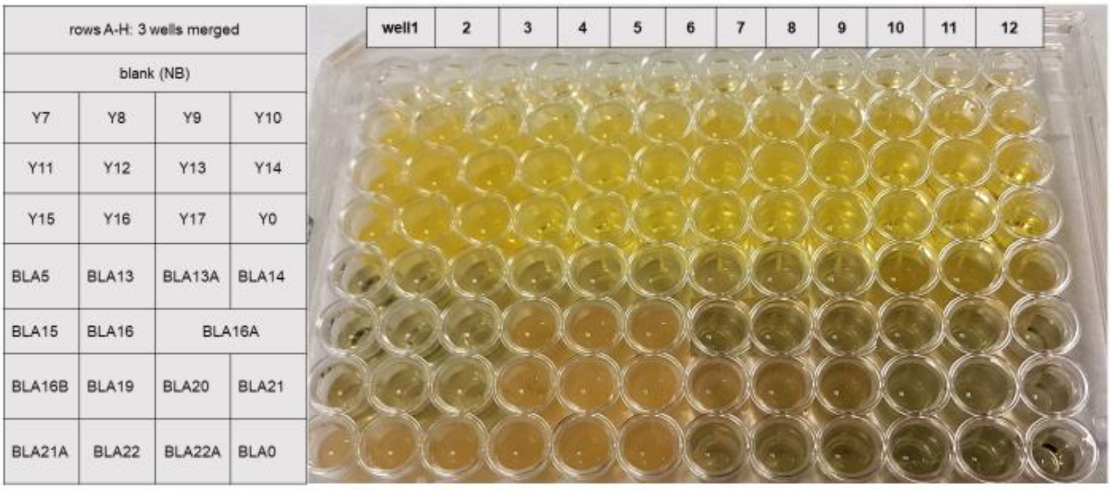
rows A-H: 3 wells merged
blank (NB)
Y7
Y8
Y9
Y10
Y11
Y12
Y13
Y14
Y15
Y16
Y17
Y0
BLA5
BLA13
BLA13A
BLA14
BLA15
BLA16
BLA16A
BLA16B
BLA19
BLA20
BLA21
BLA21A
BLA22
BLA22A
BLA0
well1

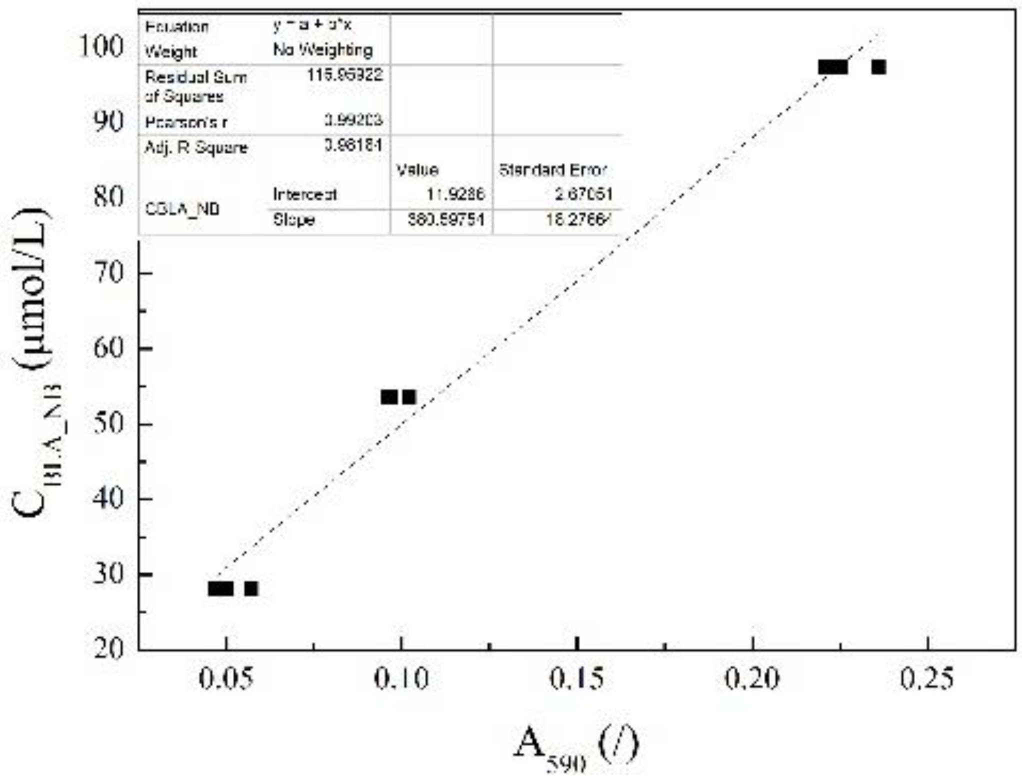
Equation
y = a + b*x
Weight
No Weighting
Residual Sum of Squares
115.95922
Pearson's r
0.99203
Adj. R Square
0.98181
Value
Standard Error
CBLA_NB
Intercept
11.9286
2.67051
Slope
380.59751
16.27664
C_BLA_NB (μmol/L)
A_590 (/)

d)

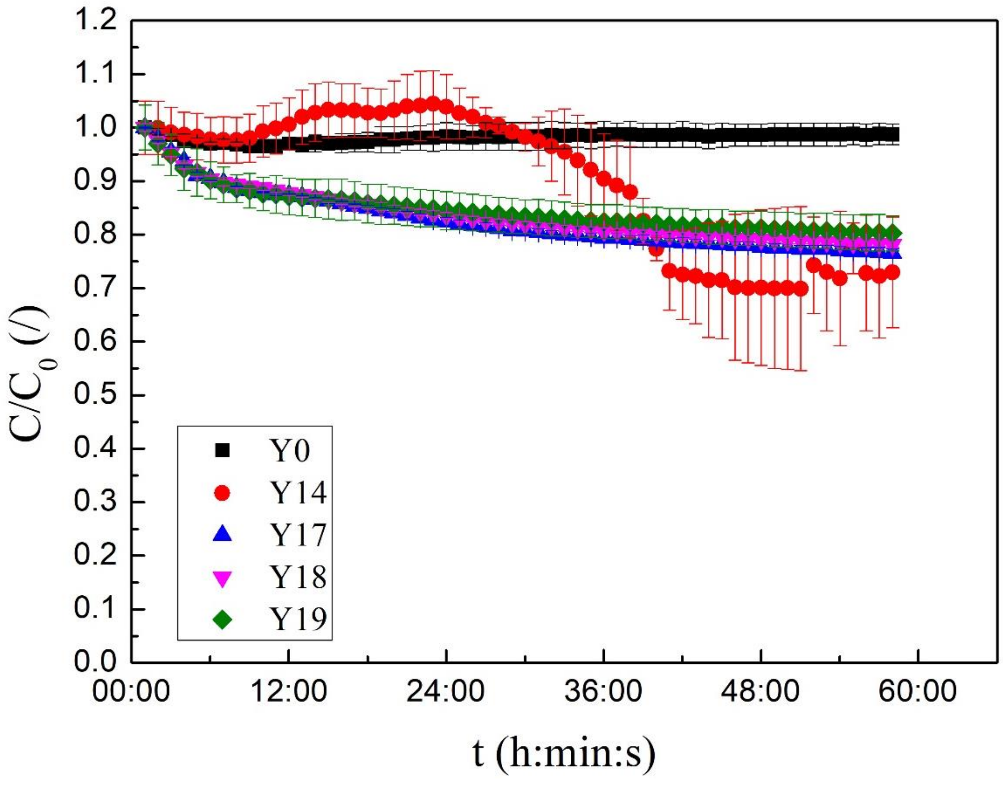
C/C_0 (/)
t (h:min:s)
Y0
Y14
Y17
Y18
Y19

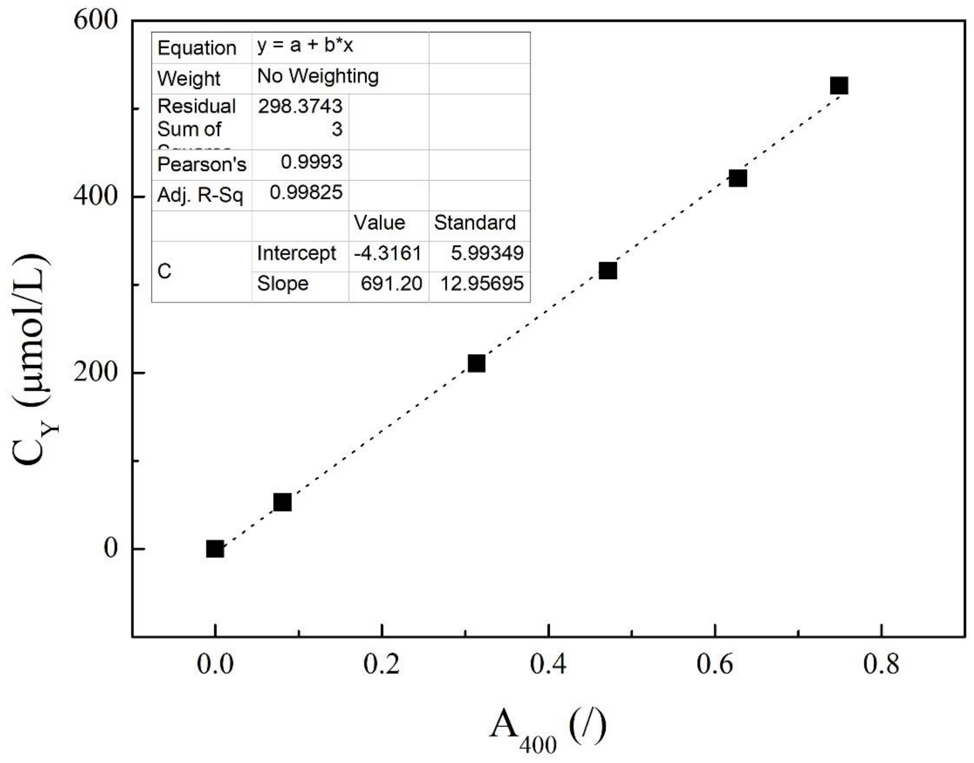

e)

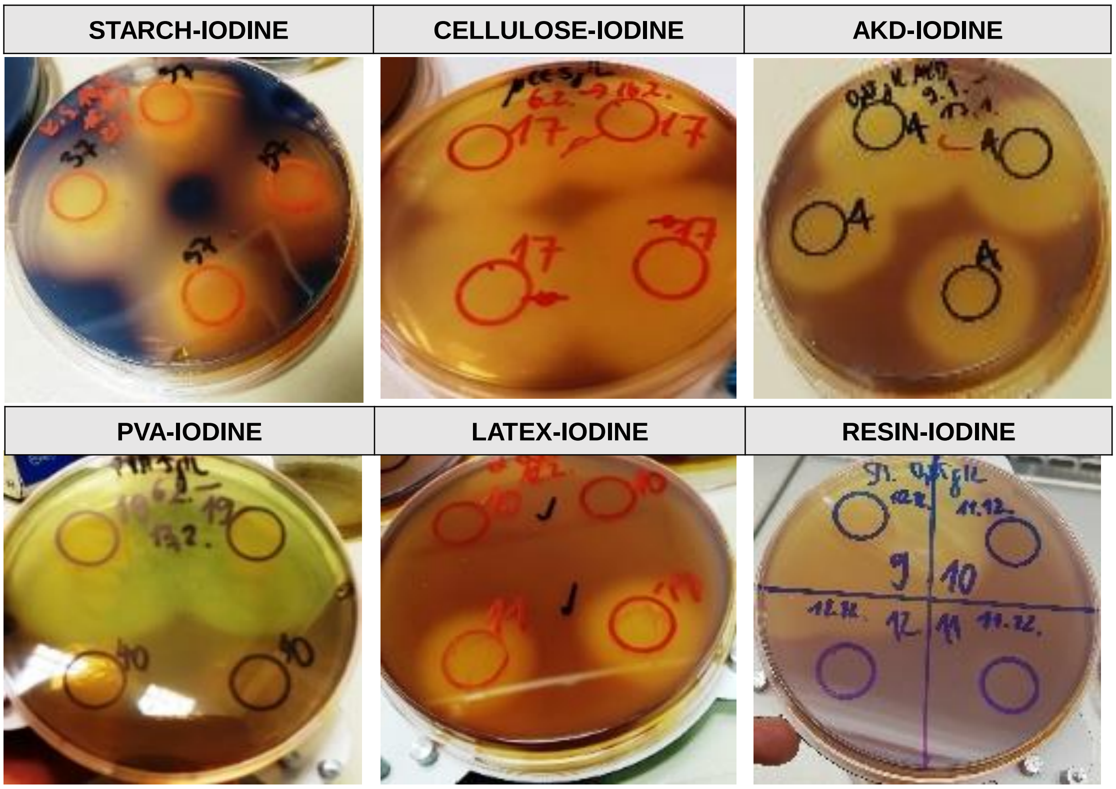

*Fig. A.1*

*Selection of isolates for usage of the carbon source that particular isolate used for isolation:* ***a)*** *BLU isolates:* ***above*** *- incubation microtiter plate with 23 BLU isolates in triplicates in NB + 5.7 mg/L blue after 66 h,* ***below*** *-* ***below*** *- calibration curve with* $R^2$*=0.995.* ***b)*** *RED isolates: above incubation microtiter plate with 12 RED isolates in NB + 15 mg/L red after 66 h in* *triplicates, below calibration curve with $R^2$=0.999.* ***c)*** *BLA isolates and unclear cometabolism test of Y isolates;* ***above*** *- microtiter plate with 11 Y and 14 BLA isolates (below) in NB +7.5 mg/L yellow or 57 mg/L black dye after 66 h in triplicates, blank value (NB).* ***Below*** *- calibration curve for black dye in NB with $R^2$=0.993.* ***d)*** *Y isolates.* ***Above*** *- kinetics of discoloration of the yellow dye by Pseudomonas sp. Y17,18,19- BF and Klebsiella sp. Y14-BTP; protocol 1. The four fastest growing Y isolates in NB, Y17,18,19 and Y14, were cleared and inoculated in M9 Y with respective ($OD_{600}$-$Y_0$): 0.03±0.1%, 0.03±0.7%, 0.02±3.9% and 0.15±6%. The values $C/C_0$ represent the absorption of cells and dye. The values of Y14 increase at the beginning due to the absorption of the cells.* ***Below*** *- the calibration curve yellow dye in M9 Y with $R^2$=0.999.* ***e)*** *CST, CE, AKD, PVA, LX and RES isolates inoculated on corresponding M9 CST, CE, AKD, PVA, LX and RES agar media. After 8 days, cell viability was checked on NB agar plates and isolates that showed a distinct discoloration after addition of Lugol were used for testing of their reservoire of carbon source usage. Photographs without corrections.* ***Above*** *from left to right: Starch-iodine complex formation and distinct discoloring around colonies of Pseudomonas sp. CST37-CF, cellulose-iodine complex and distinct discoloring around Agromyces sp. CE17-BF, AKD-iodine complex and distinct discoloring around Cellulosimicrobium sp. AKD4-BF;* ***below*** *from left to right: PVA-iodine complex and distinct discoloring around Pseudomonas sp. PVA19-CF compared to the inactivity of PVA10-BF, LX-iodine complex and distinct discoloration around Micrococcus sp. LX11- CF compared to the inactivity of LX10-BTP and RES-iodine complex formation and distinct discoloration around Pseudomonas sp. RES9-CF compared to the inactivity of RES10-BTP, RES11-BTP, RES12-BTP.*

a)

| rows A-H: 3 wells merged | | |
|---|---|---|
| blank (100 µl) + lugol (100 µl) | blank (100 µl) | |
| RES16 | RES19 | ST9 |
| ST12 | ST14 | ST16 |
| CST10A | CST10B | CST10C |
| CST16 | CST37 | CST5A |
| CST5B | CST5C | CE12 |
| CE17 | CE21 | CE7 |
| NST20 | NST22 | NST22A |

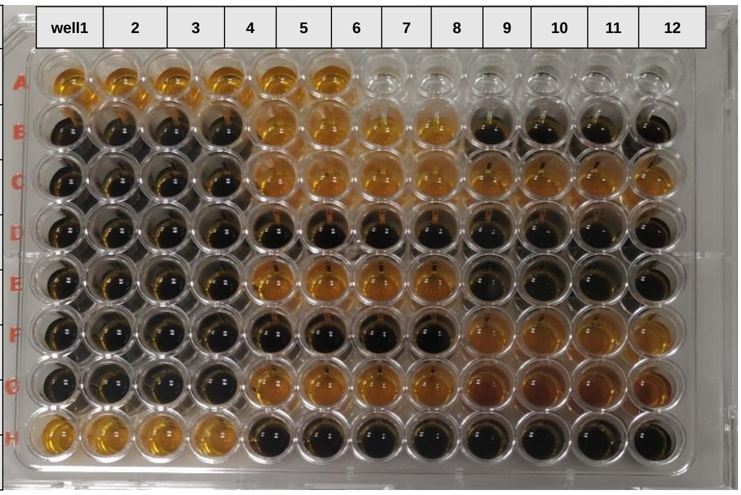

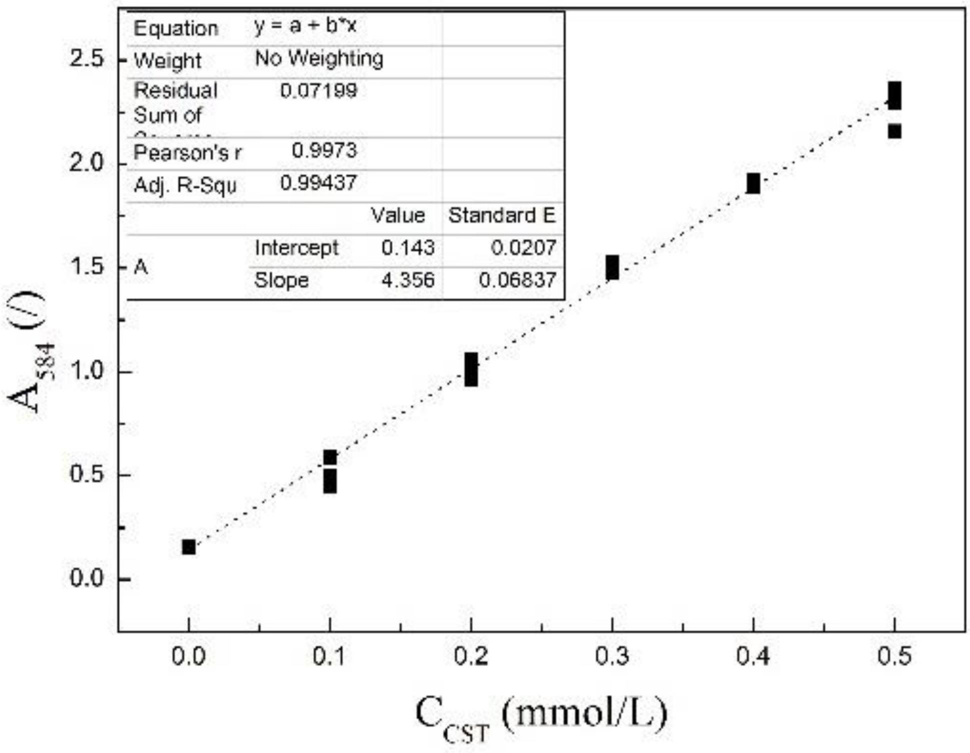

b)

| rows A-H | | |
|---|---|---|
| blank (100 µl) + lugol (100 µl) | blank (100 µl) | |
| RES16 | RES19 | ST9 |
| ST12 | ST14 | ST16 |
| CST10A | CST10B | CST10C |
| CST16 | CST37 | CST5A |
| CST5B | CST5C | CE12 |
| CE17 | CE21 | CE7 |
| NST20 | NST22 | NST22A |

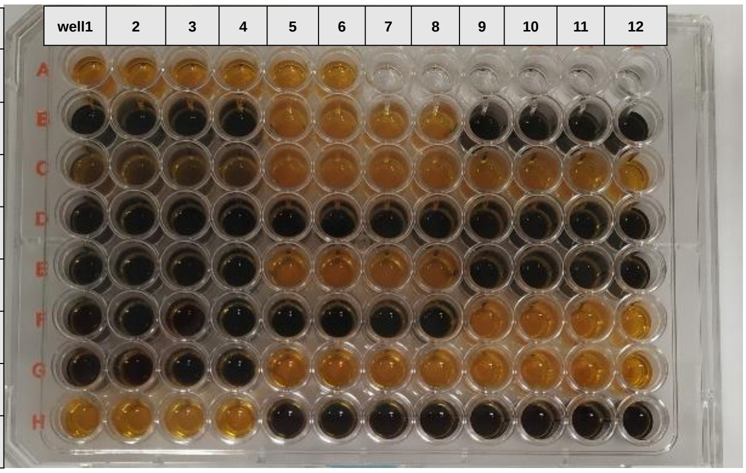

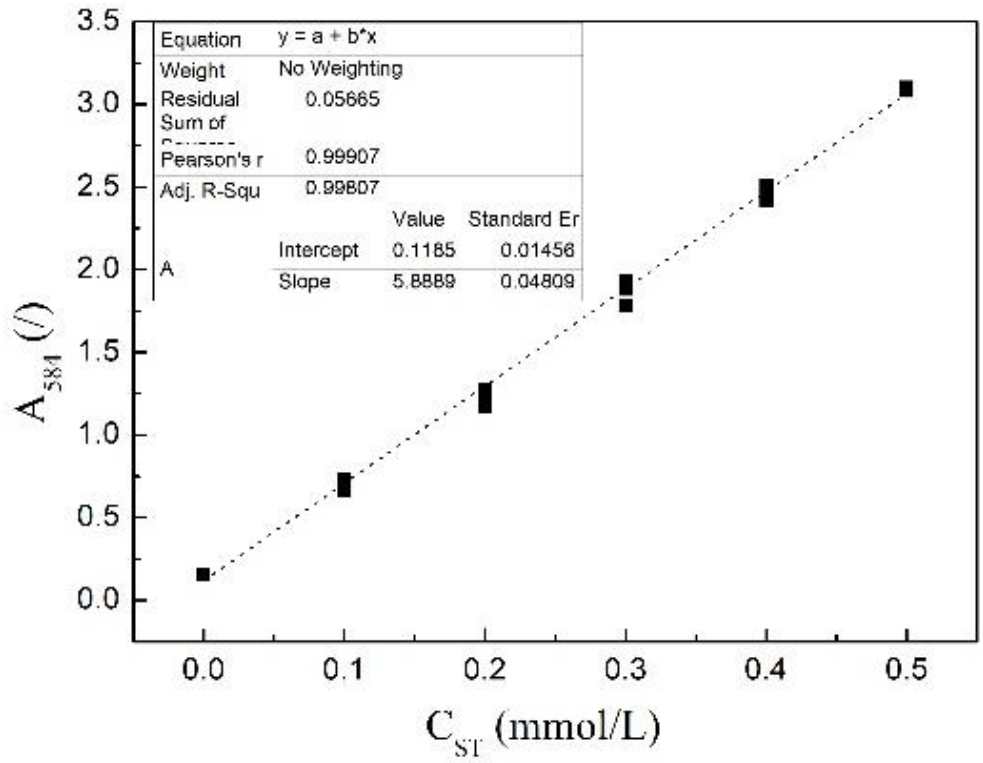

c)

| rows A-H: 3 wells merged | | |
|---|---|---|
| blank (100 µl) + lugol (100 µl) | control (100 µl) + lugol (100 µl) | |
| RES16 | RES19 | ST9 |
| ST12 | ST14 | ST16 |
| CST10A | CST10B | CST10C |
| CST16 | CST37 | CST5A |
| CST5B | CST5C | CST37 |
| CE17 | CE21 | CE7 |
| NST20 | NST22 | NST22A |

| well1 | 2 | 3 | 4 | 5 | 6 | 7 | 8 | 9 | 10 | 11 | 12 |
|---|---|---|---|---|---|---|---|---|---|---|---|

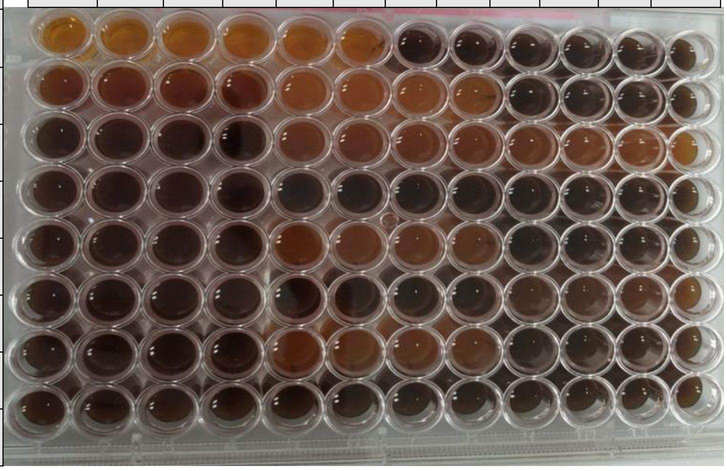

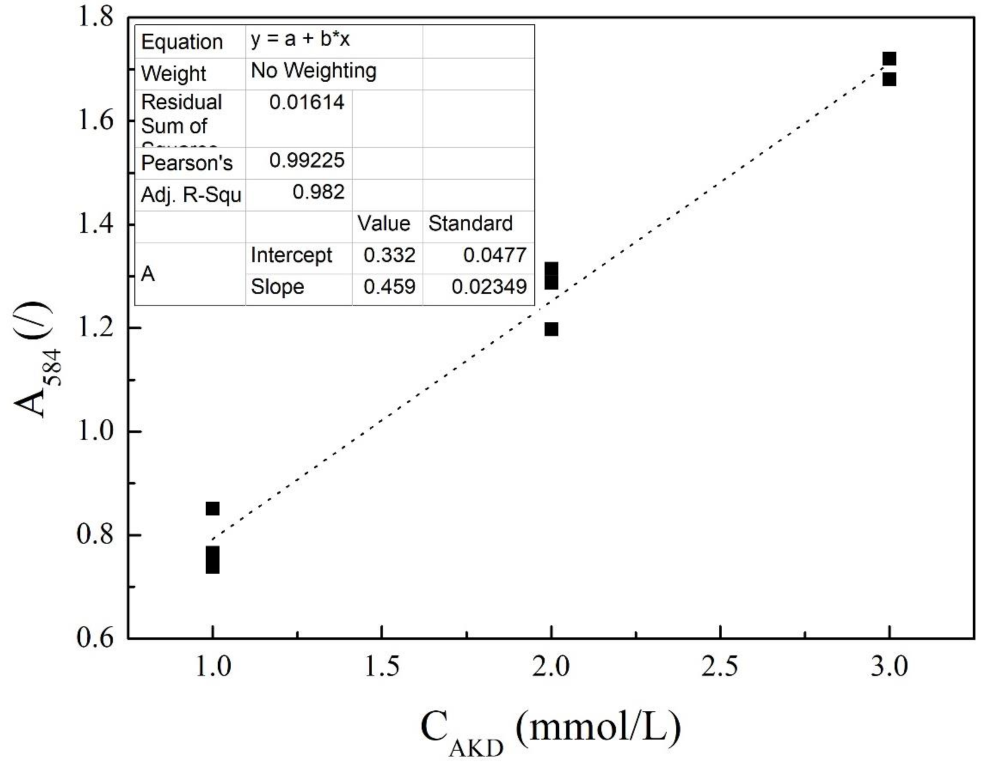

d)

| rows A-H | | |
|---|---|---|
| blank (200 µl) | control (200 µl) | |
| blank | RED15A | RED16B |
| AKD4 | AKD4A | AKD5 |
| ALD5A | AKD13A | AKD19 |
| PVA19 | PVA20 | PVA3 |
| PVA7 | PVA14 | LX8 |
| LX11 | LX13 | RES9 |
| RES13 | RES15 | AKD13 |

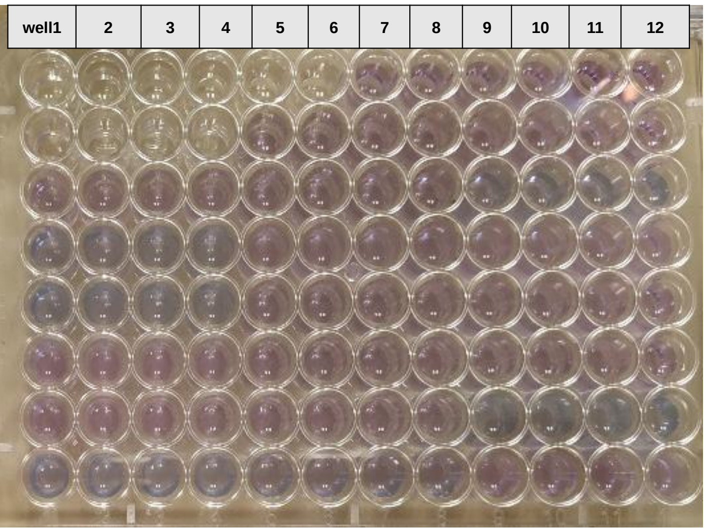

| rows A-H: 3 wells merged | | | |
|---|---|---|---|
| blank (M9) | | | |
| control | | AKD5A | AKD13A |
| PVA19 | AKD19 | PVA20 | PVA3 |
| PVA7 | PVA14 | LX8 | LX11 |
| LX13 | RES9 | RES13 | RES15 |
| AKD13 | RES19 | RES16 | ST9 |
| ST12 | ST14 | ST16 | CST10A |
| CST10B | CST10C | CST16 | CST37 |

| well1 | 2 | 3 | 4 | 5 | 6 | 7 | 8 | 9 | 10 | 11 | 12 |
|---|---|---|---|---|---|---|---|---|---|---|---|

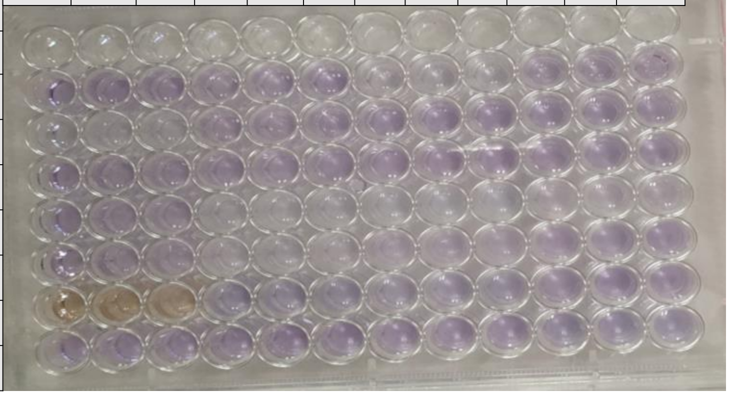

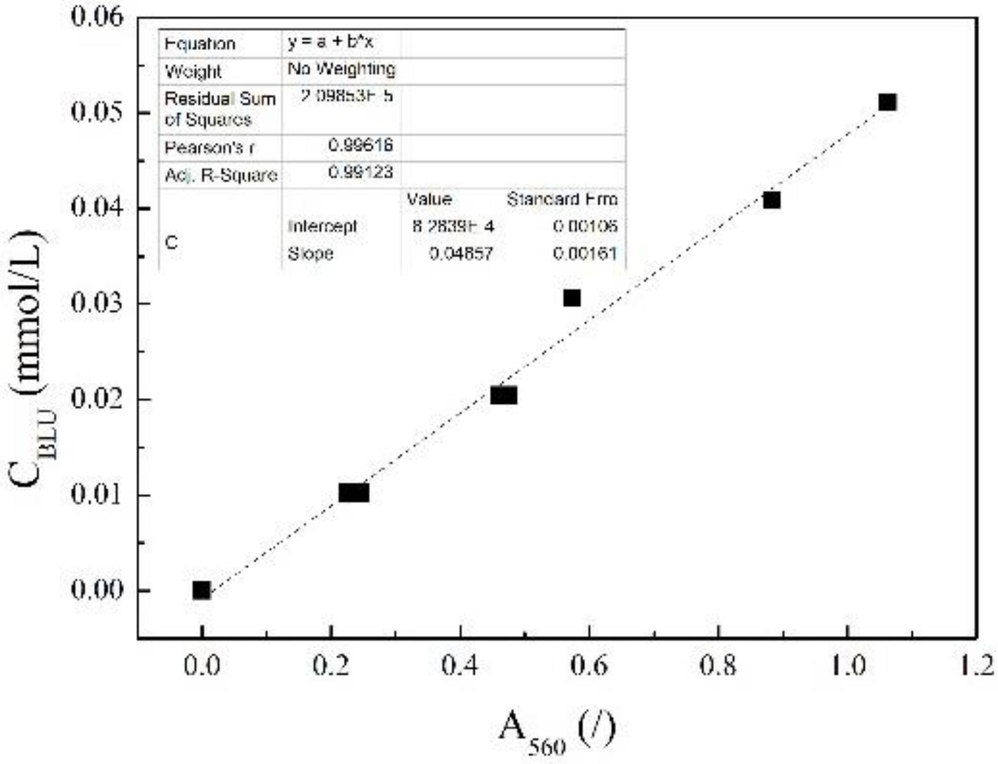

e)

| rows A-H | | |
|---|---|---|
| blank (200 µl) | control (200 µl) | |
| blank | | |
| AKD4 | AKD4A | AKD5 |
| ALD5A | AKD13A | AKD19 |
| PVA19 | PVA20 | PVA3 |
| PVA7 | PVA14 | LX8 |
| LX11 | LX13 | RES9 |
| RES13 | RES15 | AKD13 |

| well1 | 2 | 3 | 4 | 5 | 6 | 7 | 8 | 9 | 10 | 11 | 12 |
|---|---|---|---|---|---|---|---|---|---|---|---|

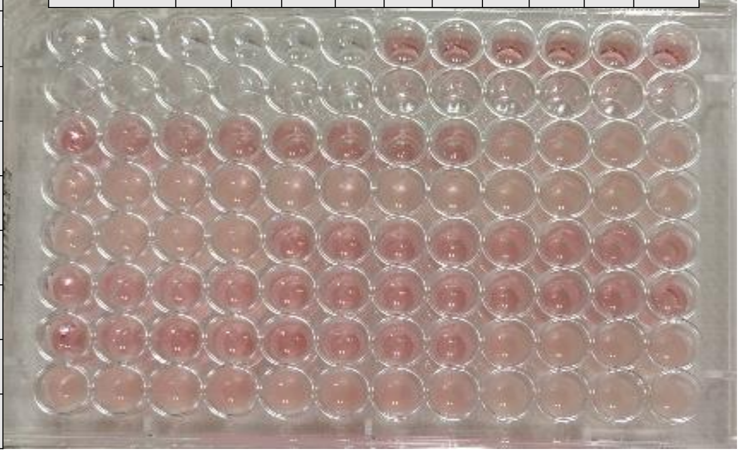

| | | | |
|---|---|---|---|
| rows A-H: 3 wells merged | | | |
| blank (M9) | | | |
| control | | AKD5A | AKD13A |
| PVA19 | AKD19 | PVA20 | PVA3 |
| PVA7 | PVA14 | LX8 | LX11 |
| LX13 | RES9 | RES13 | RES15 |
| AKD13 | RES16 | RES19 | ST9 |
| ST12 | ST14 | ST16 | CST10A |
| CST10B | CST10C | CST16 | CST37 |

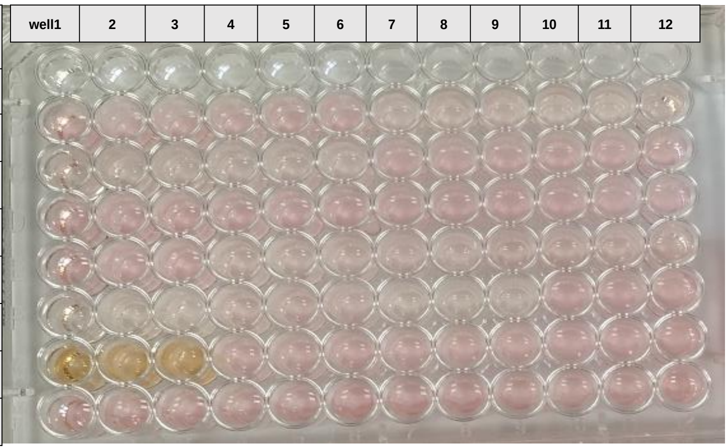

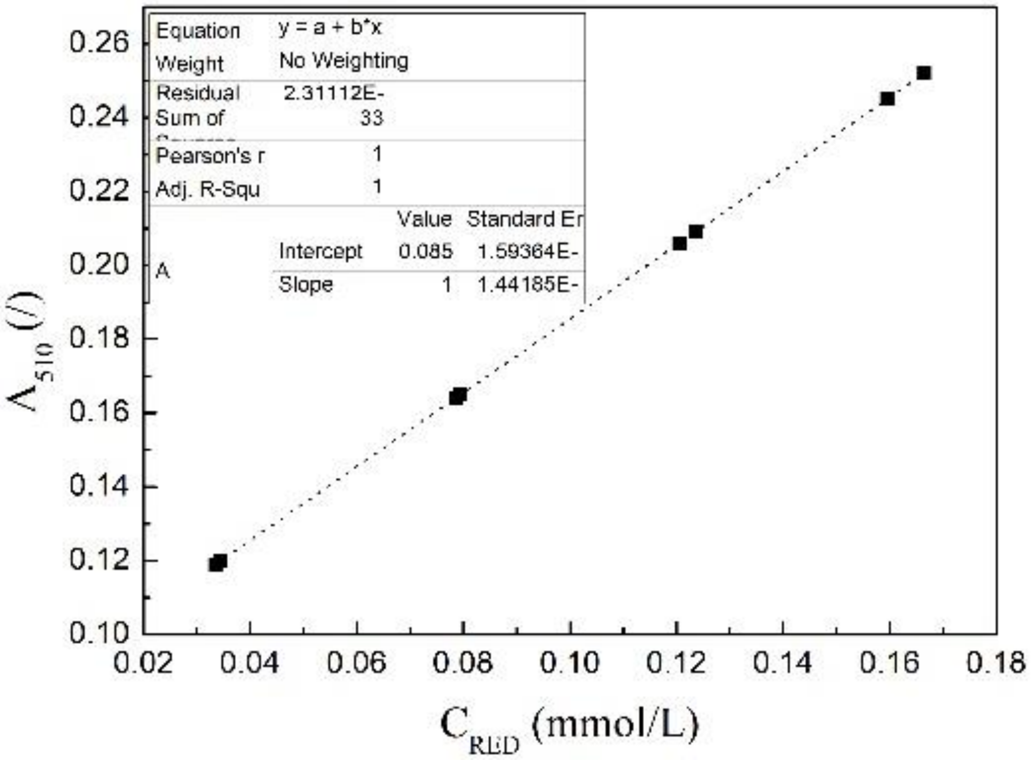

f)

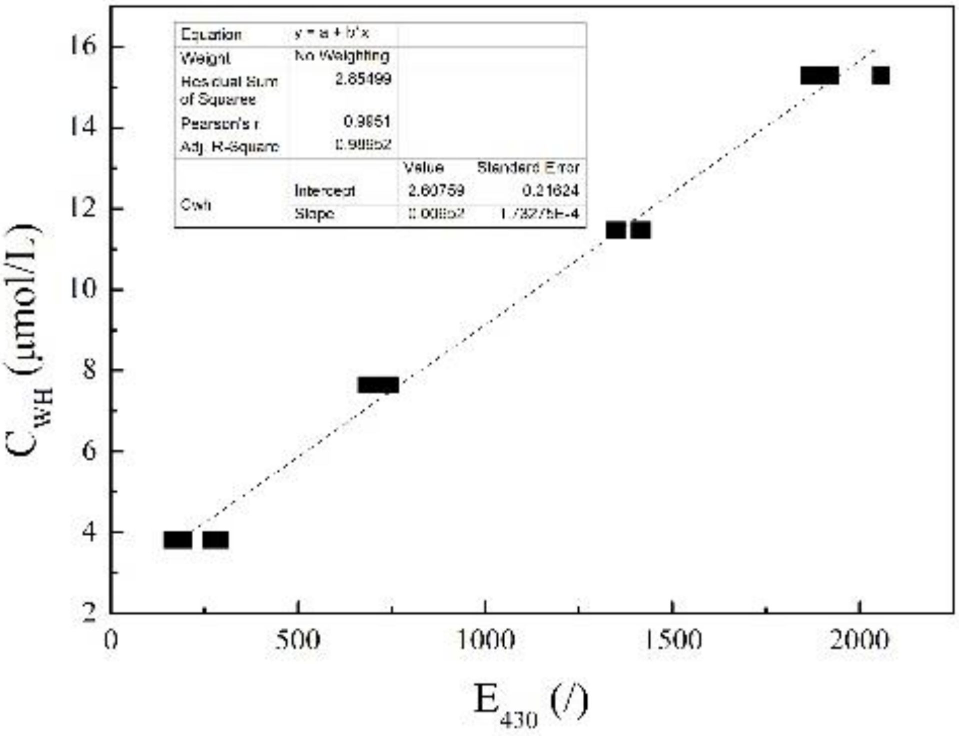

g)

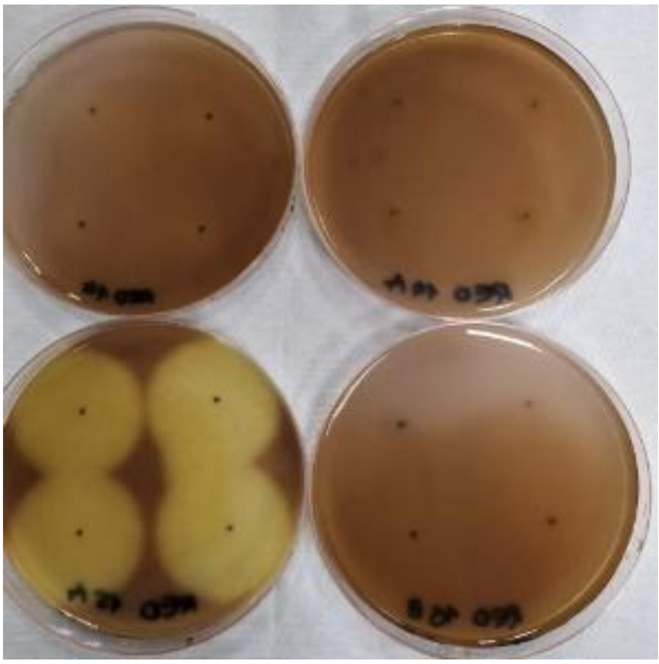

*Fig. A.2*

*Tests for reservoire of carbon source usage.* ***a)*** *CST:* ***Above*** *- microtiter plate after addition of Lugol. Bacteria incubated in 0.5 g/L CST in M9, in a sterile microtiter plate with flat bottom, 200 µL per well, in quadruplicates. After 4 days at 25°C 100 µL contents were transferred into new microtiter plate, absorption at $\lambda_{584}$ was recorded before and after addition of 100 µL Lugol; blank: M9 salts, control: sterile CST media + Lugol, sample: CST media with bacteria*

*+ Lugol.* ***Below*** *- calibration curve ($R^2$=0.997) under identical conditions: 100 μL Lugol + 100 μL CST media, $\lambda_{584}$.* ***b)*** *ST.* ***Above*** *- microtiter plate after addition of Lugol,* ***below*** *- calibration curve ($R^2$=0.999).* ***c)*** *AKD:* ***Above*** *- microtiter plate after addition of Lugol in 4 g/L AKD,* ***below*** *- calibration curve ($R^2$=0.999).* ***d)*** *BLU in M9 Glc: Incubation was performed in a sterile flat microtiter plate containing 200 μL (4.7 mg/L BLU in M9 Glc).* ***Above*** *- microtiter plate after 4 days at 25°C when $OD_{600}$ was recorded; blank value: M9 salts, control: sterile medium; bacteria incubated in quadruplicates. After 4 days at 25°C contents of wells were emptied into sterile microtubes, after 5 minutes at 5 rcf - 150 μL supernatant in triplicates transferred in new microtiter plate and $\lambda_{max}$ recorded. Example of the supernatants in the* ***middle****.* ***Below*** *is the calibration curve ($R^2$=0.996) from which the unutilized concentration of the dye was calculated.* ***e)*** *RED in M9 Glc* ***Above*** *- incubation microtiter plate containing 200 μL (4.7 mg/L RED in M9 Glc) after 5 days at 25°C,* ***middle*** *- supernatants,* ***below*** *- calibration curve ($R^2$=0.999).* ***f)*** *WH in M9 Glc. Incubated in transparent microtiter plate with 200 μL content (4.7 mg/L WH in M9 Glc). Content was transferred into sterile microtubes after 4 days at 25°C and after 5 min at 5 rcf - 150 μL supernatants transferred in triplicates into black microtiter plates and fluorescence emission at 440 nm recorded after excitation at $\lambda_{315}$. Calibration curve ($R^2$=0.996).* ***g)*** *Cellulose- iodine complex formation after addition of Lugol and distinct discoloration around the cellulolytic activity of Agromyces sp. RED15A- BF compared to the inactivity of Stenotrophomonas sp. NV -RED14- BF, Micrococcus sp. NV -RED14A- BF and Stenotrophomonas sp. NV -RED16B- BF.*

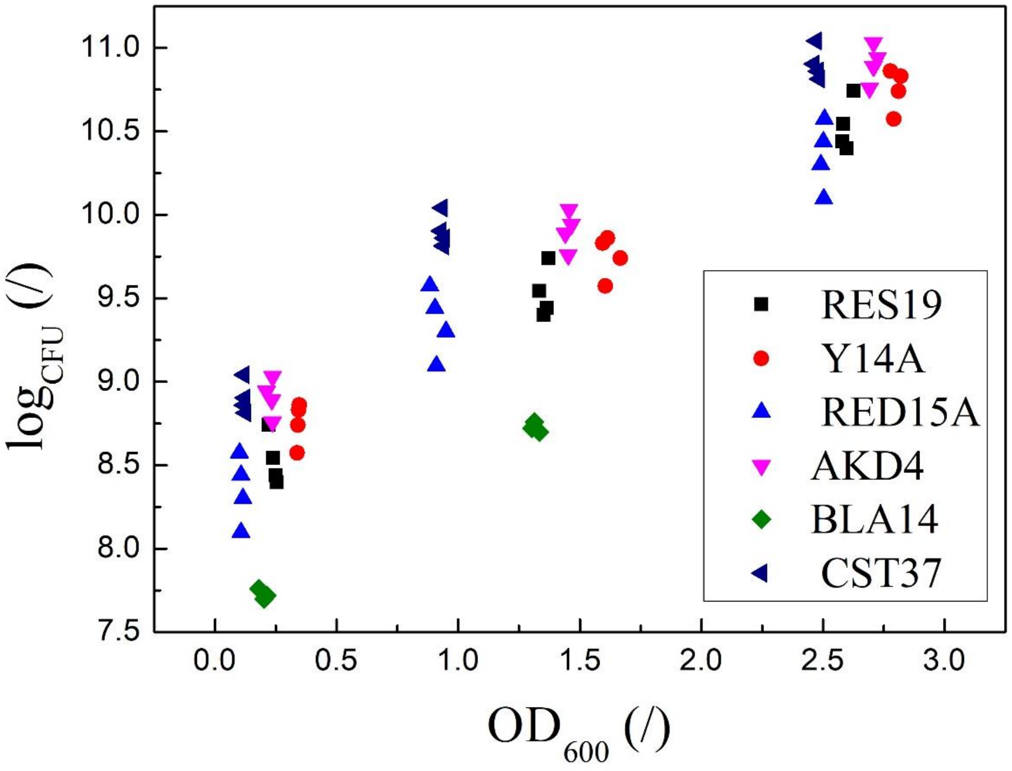

*Fig. A.3*

*Calibration of the growth of the 6 selected bacteria - CFU vs. $OD_{600}$.*

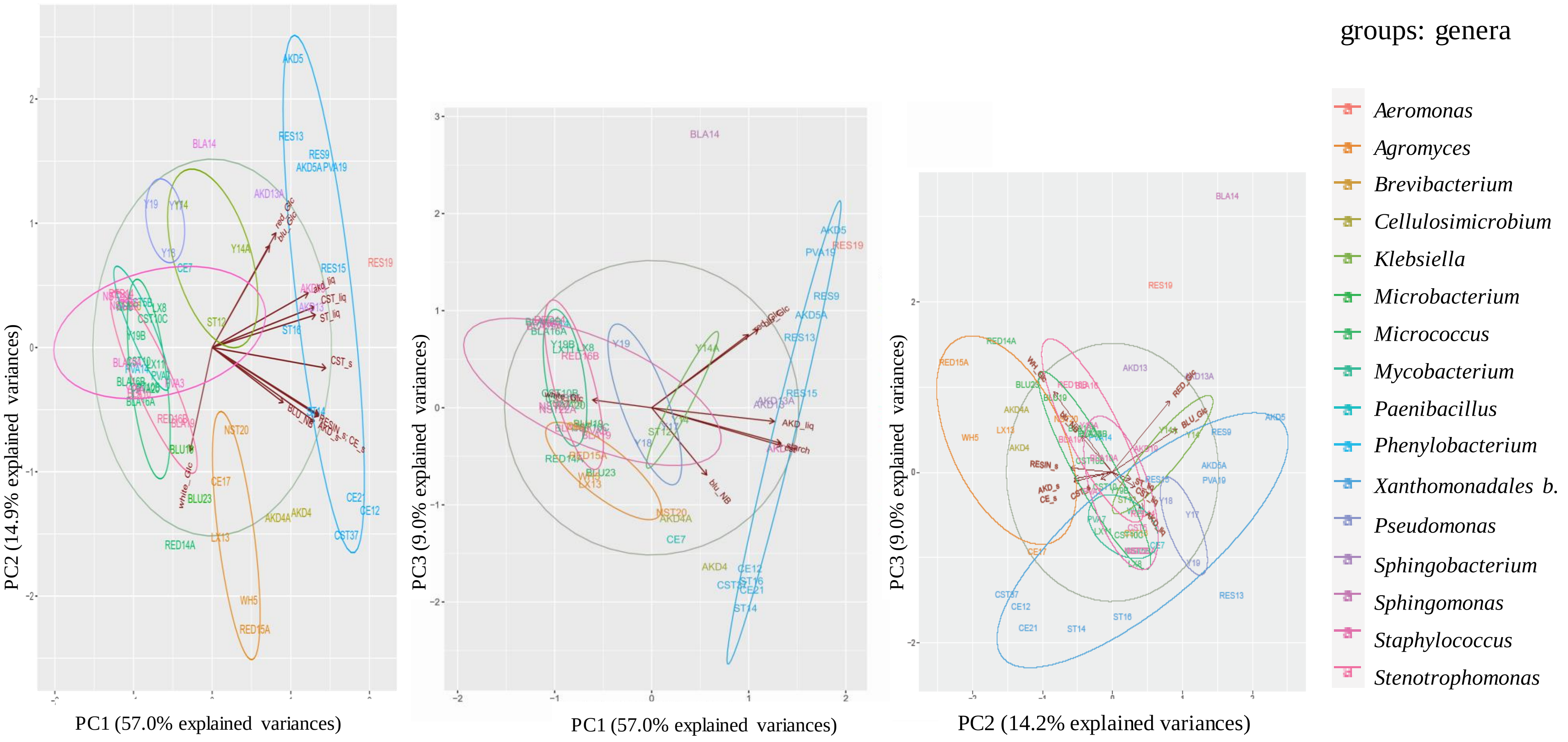

*Fig. A.4*

*PCA of tests for the repertoire of carbon sources of 58 isolates in liquid and solid media with genus clusters. PC1PC2 (**left**), PC1PC3 (**middle**), PC2PC3 (**right**).*

a)

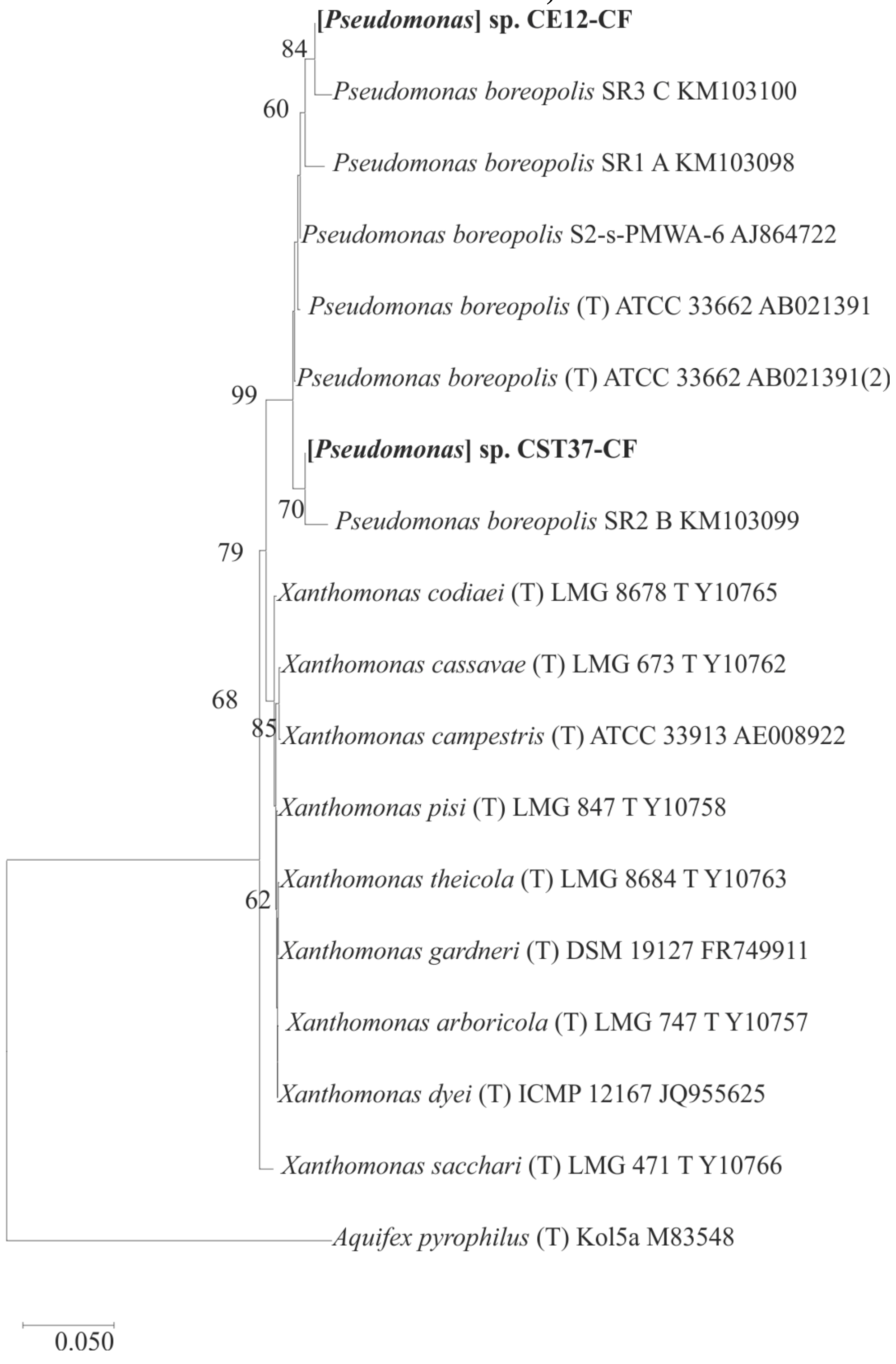

**[*Pseudomonas*] sp. CE12-CF**
84
*Pseudomonas boreopolis* SR3 C KM103100
60
*Pseudomonas boreopolis* SR1 A KM103098
*Pseudomonas boreopolis* S2-s-PMWA-6 AJ864722
*Pseudomonas boreopolis* (T) ATCC 33662 AB021391
*Pseudomonas boreopolis* (T) ATCC 33662 AB021391(2)
99
**[*Pseudomonas*] sp. CST37-CF**
70
*Pseudomonas boreopolis* SR2 B KM103099
79
*Xanthomonas codiaei* (T) LMG 8678 T Y10765
*Xanthomonas cassavae* (T) LMG 673 T Y10762
68
85
*Xanthomonas campestris* (T) ATCC 33913 AE008922
*Xanthomonas pisi* (T) LMG 847 T Y10758
*Xanthomonas theicola* (T) LMG 8684 T Y10763
62
*Xanthomonas gardneri* (T) DSM 19127 FR749911
*Xanthomonas arboricola* (T) LMG 747 T Y10757
*Xanthomonas dyei* (T) ICMP 12167 JQ955625
*Xanthomonas sacchari* (T) LMG 471 T Y10766
*Aquifex pyrophilus* (T) Kol5a M83548
0.050

b)

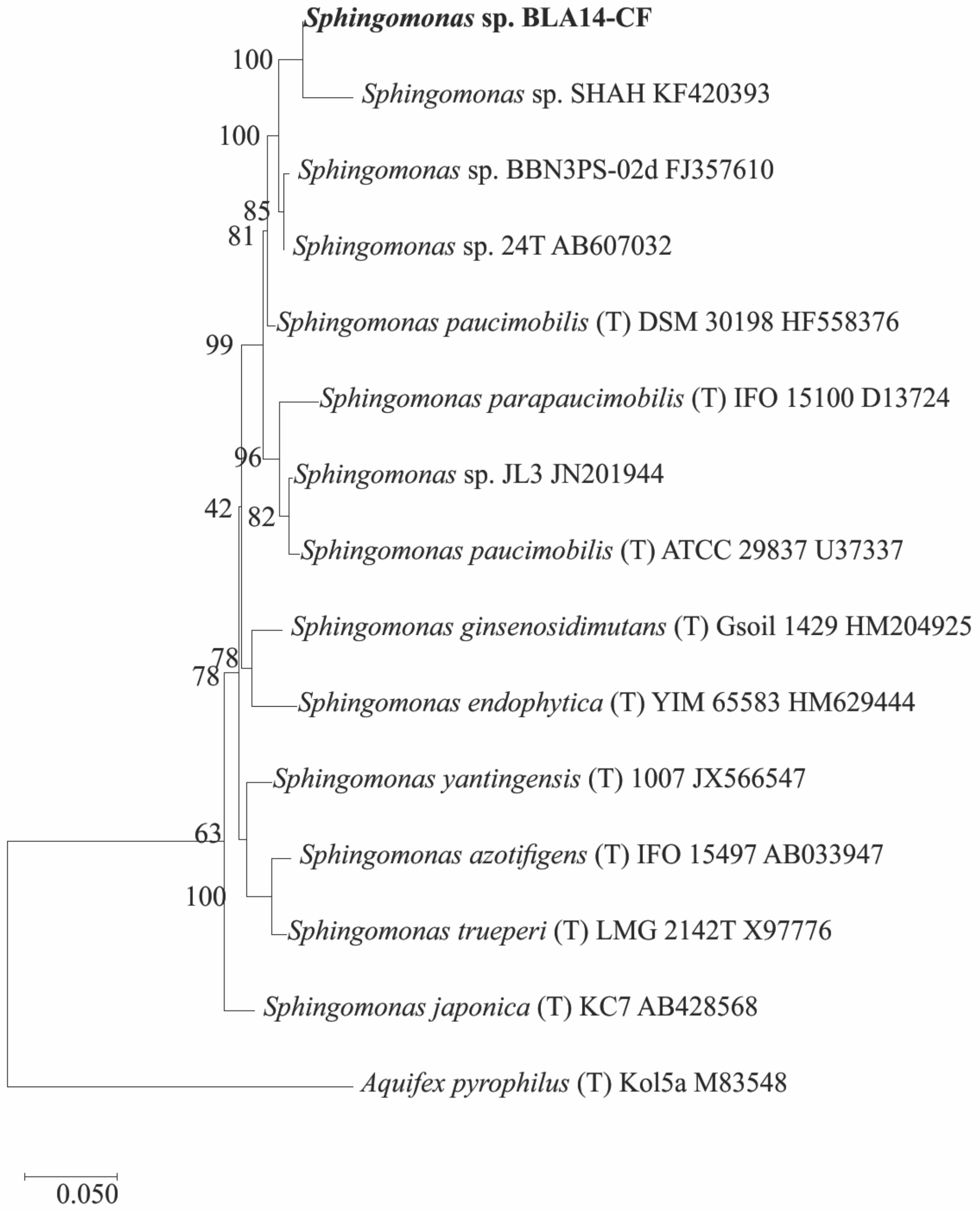
Sphingomonas sp. BLA14-CF
Sphingomonas sp. SHAH KF420393
Sphingomonas sp. BBN3PS-02d FJ357610
Sphingomonas sp. 24T AB607032
Sphingomonas paucimobilis (T) DSM 30198 HF558376
Sphingomonas parapaucimobilis (T) IFO 15100 D13724
Sphingomonas sp. JL3 JN201944
Sphingomonas paucimobilis (T) ATCC 29837 U37337
Sphingomonas ginsenosidimutans (T) Gsoil 1429 HM204925
Sphingomonas endophytica (T) YIM 65583 HM629444
Sphingomonas yantingensis (T) 1007 JX566547
Sphingomonas azotifigens (T) IFO 15497 AB033947
Sphingomonas trueperi (T) LMG 2142T X97776
Sphingomonas japonica (T) KC7 AB428568
Aquifex pyrophilus (T) Kol5a M83548
100
100
85
81
99
96
42
82
78
78
63
100
0.050

c)

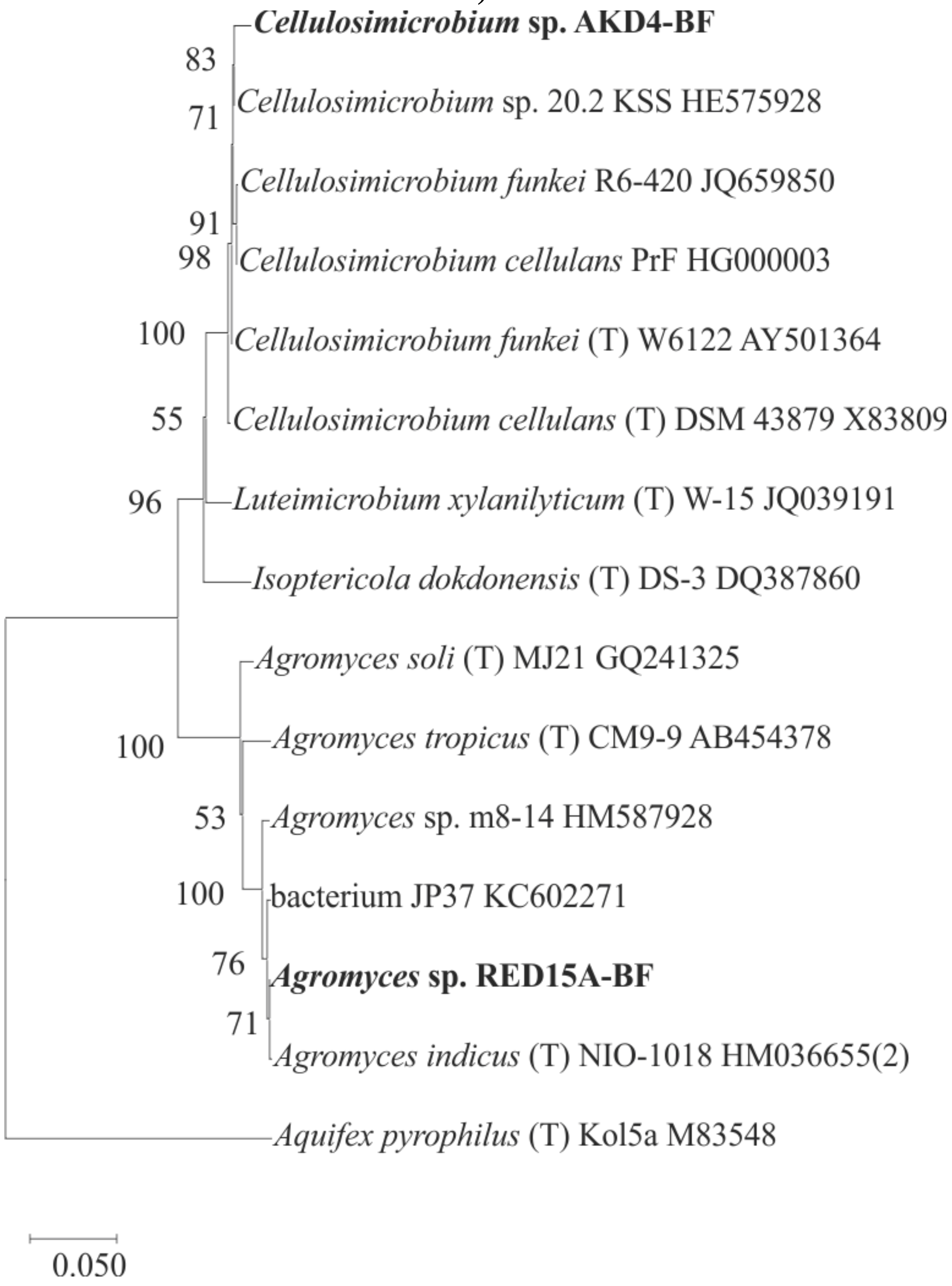
Cellulosimicrobium sp. AKD4-BF
Cellulosimicrobium sp. 20.2 KSS HE575928
Cellulosimicrobium funkei R6-420 JQ659850
Cellulosimicrobium cellulans PrF HG000003
Cellulosimicrobium funkei (T) W6122 AY501364
Cellulosimicrobium cellulans (T) DSM 43879 X83809
Luteimicrobium xylanilyticum (T) W-15 JQ039191
Isoptericola dokdonensis (T) DS-3 DQ387860
Agromyces soli (T) MJ21 GQ241325
Agromyces tropicus (T) CM9-9 AB454378
Agromyces sp. m8-14 HM587928
bacterium JP37 KC602271
Agromyces sp. RED15A-BF
Agromyces indicus (T) NIO-1018 HM036655(2)
Aquifex pyrophilus (T) Kol5a M83548
83
71
91
98
100
55
96
100
53
100
76
71
0.050

d)

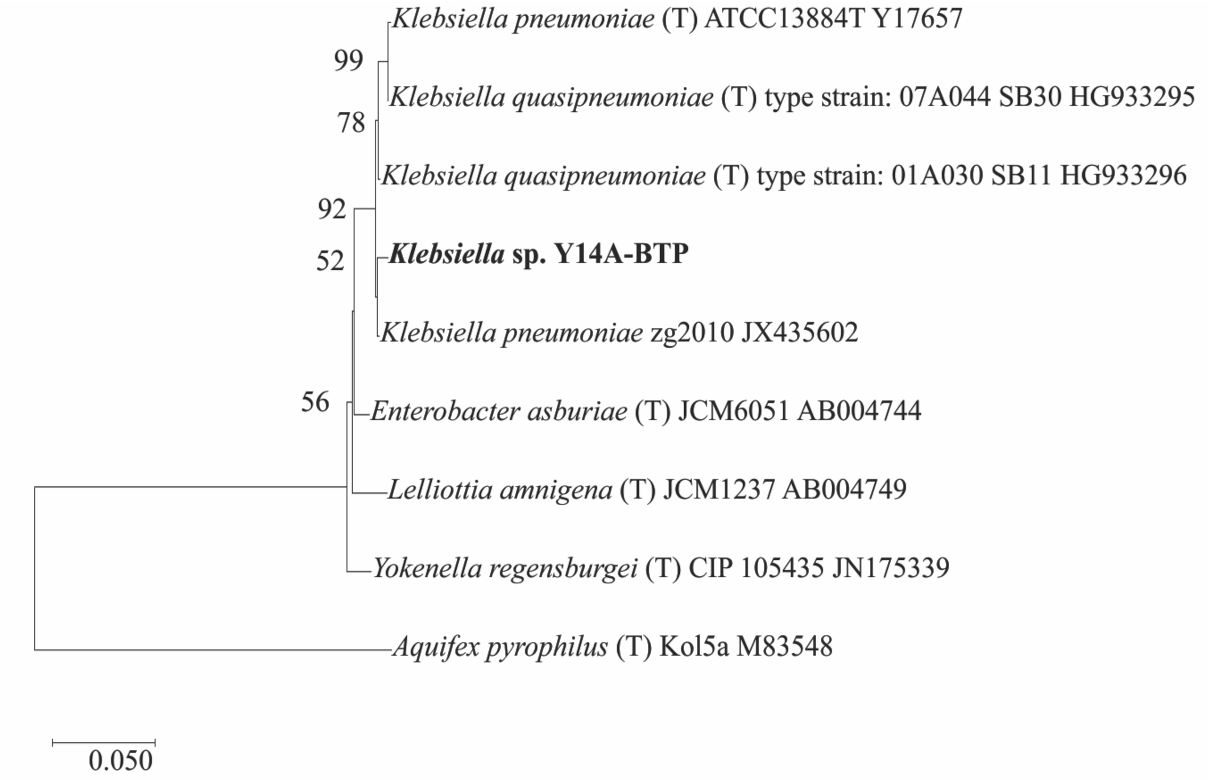

*Fig. A.5*

*Detailed phylogenetic trees of the six selected isolates with their closest neighbors are presented. Aquifex pyrophilus (T) was used as outlier. The evolutionary history was derived using the Neighbor-Joining method (Saitou and Nei 1987).* ***a)*** *Xanthomonadales bacterium sp. CST37 together with CE12,* ***b)*** *Sphingomonas sp. BLA14-CF,* ***c)*** *Cellulosimicrobium sp. AKD4-BF together with Agromyces sp. RED15A and* ***d)*** *Klebsiella sp. Y14A-BTP.*

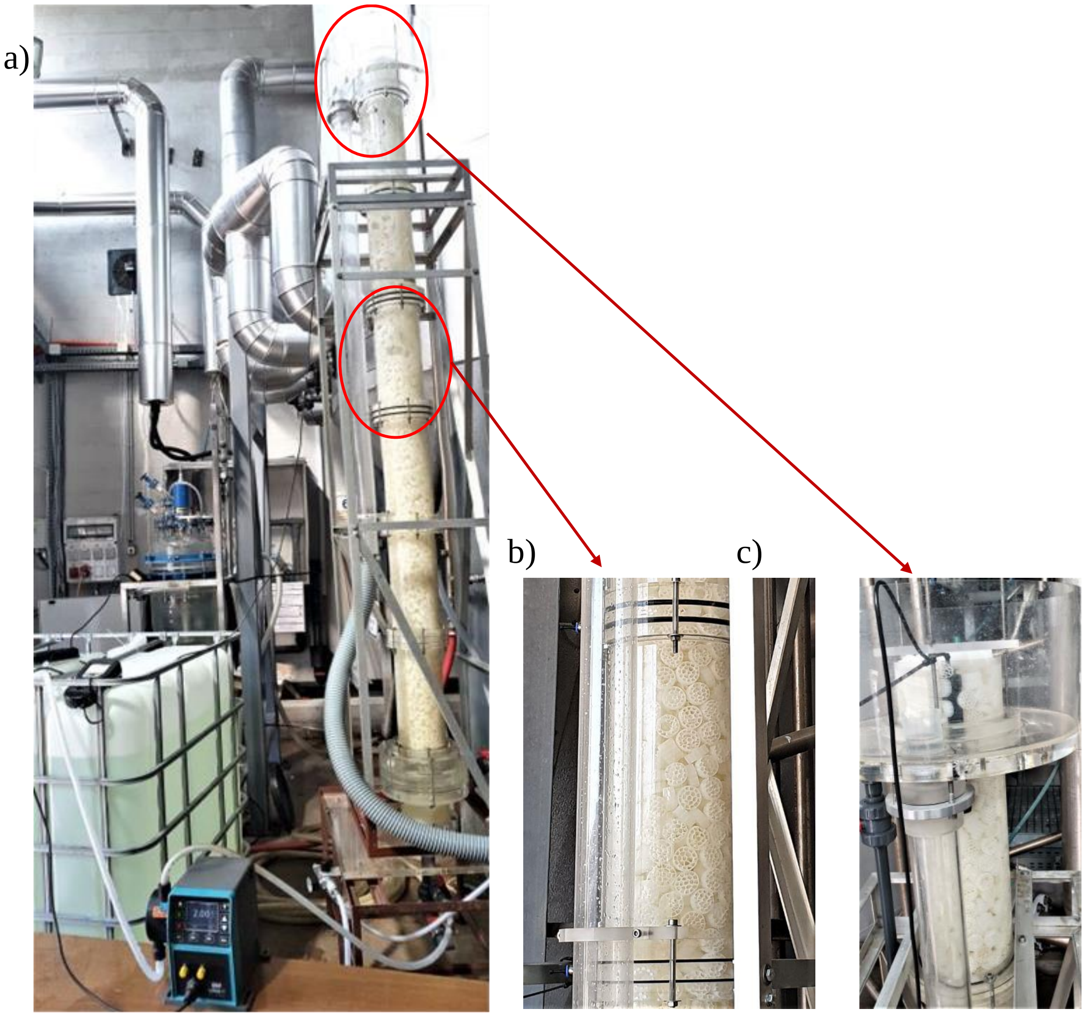
a)
b)
c)

d)

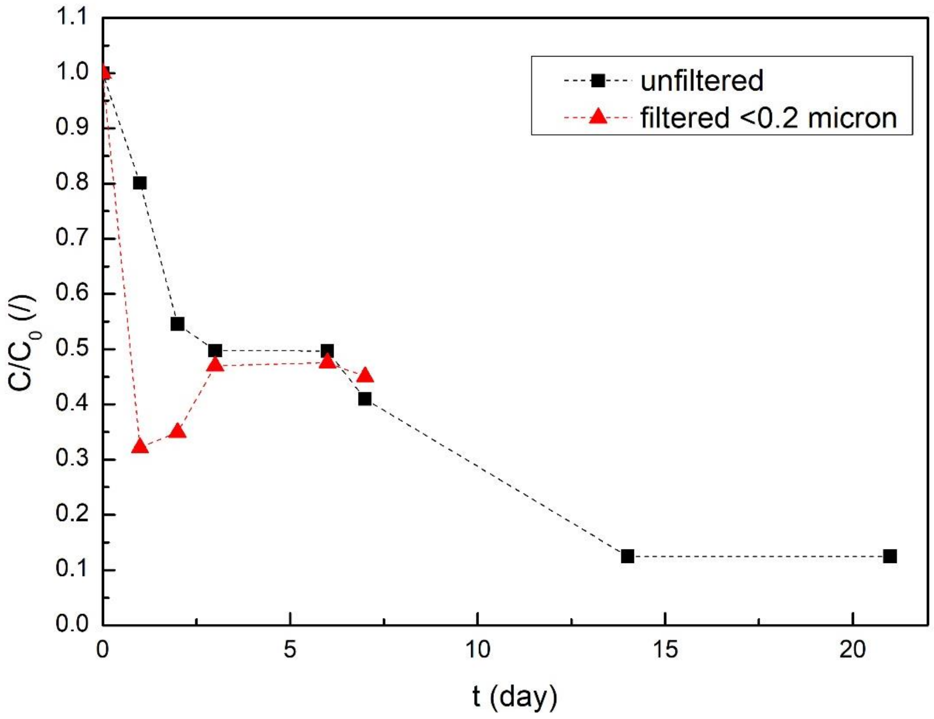

e)

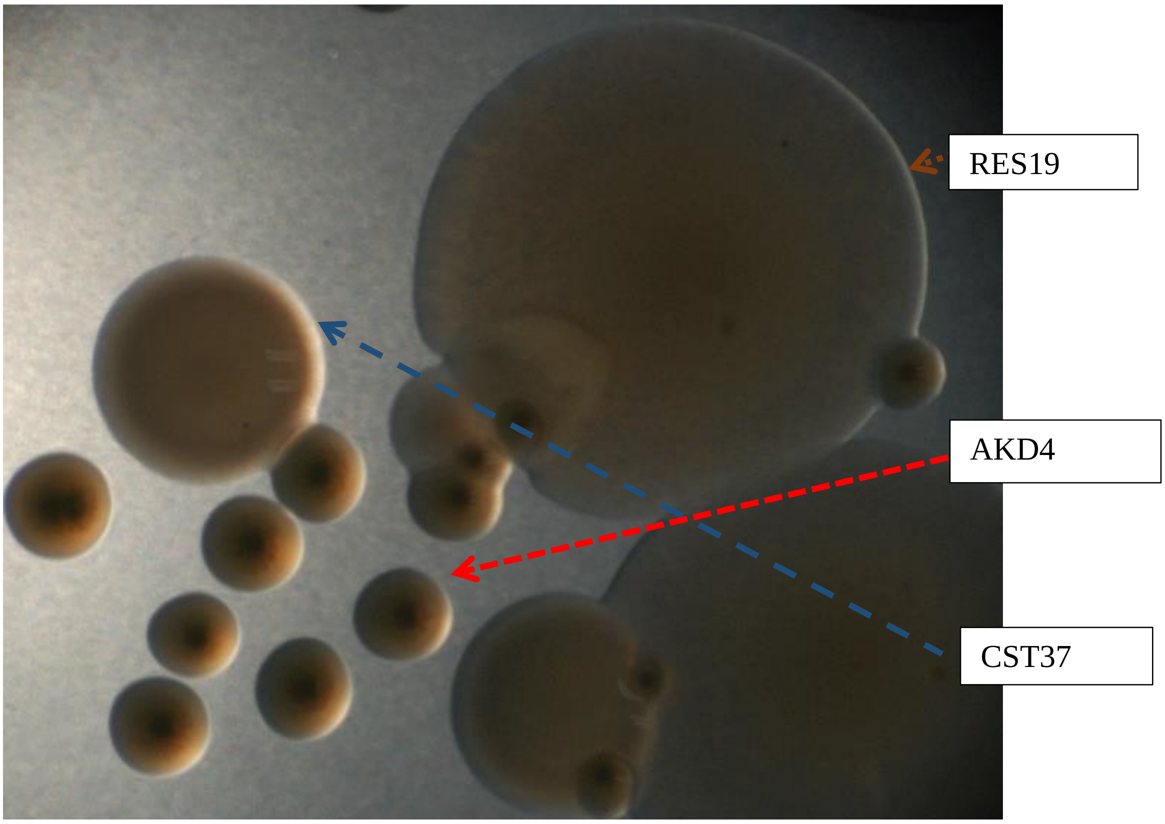

*Fig. A.6*

*Pilot test (for proof of concept only):* ***a)*** *Photograph of the set-up: a 33 L column with parallel flow of whitewater and air entering at the bottom of the column by means of Watson Marlow qdos 30d pump;* ***b)*** *part of the column filled with plastic carriers "Kaldens". Mixing* *of the carriers almost absent during the experiment;* ***c)*** *upper part of the column with inserted dissolved oxygen electrode;* ***d)*** *$C/C_0$ from COD measurements of unfiltered and filtered through 0.2 μm membrane influent and effluent whitewater as a function of time. On the 8$^{th}$ day of the experiment a recycling ratio of 1 was established;* ***e)*** *after 7 days of the experiment, carriers 20 cm below the upper part of the column were transferred into sterile centrifuge tubes with 20 mL of 9 g/L NaCl, homogenized, diluted and incubated on NB agar. A representative image of the colonies was obtained with the microscope Motic SMZ-168- BL in a parfocal 6.7:1 zoom. The cfu count was as follows: 3-x RES19, 2-x CST37 and 13-x AKD4.*
